# Neuronal mechanisms of novelty seeking

**DOI:** 10.1101/2021.03.12.435019

**Authors:** Takaya Ogasawara, Fatih Sogukpinar, Kaining Zhang, Yang-Yang Feng, Julia Pai, Ahmad Jezzini, Ilya E. Monosov

## Abstract

Humans and other primates interact with the world by observing and exploring visual objects. In particular, they often seek out the opportunities to view novel objects that they have never seen before, even when they have no extrinsic primary reward value. However, despite the importance of novel visual objects in our daily life, we currently lack an understanding of how primate brain circuits control the motivation to seek out novelty. We found that novelty-seeking is regulated by a small understudied subcortical region, the zona incerta (ZI). In a task in which monkeys made eye movements to familiar objects to obtain the opportunity to view novel objects, many ZI neurons were preferentially activated by predictions of future novel objects and displayed burst excitations before gaze shifts to gain access to novel objects. Low intensity electrical stimulation of ZI facilitated gaze shifts, while inactivations of ZI reduced novelty-seeking. Surprisingly, additional experiments showed that this ZI-dependent novelty seeking behavior is not regulated by canonical neural circuitry for reward seeking. The habenula-dopamine pathway, known to reflect reward predictions that control reward seeking, was relatively inactive during novelty-seeking behavior in which novelty had no extrinsic reward value. Instead, high channel-count electrophysiological experiments and anatomical tracing identified a prominent source of control signals for novelty seeking in the anterior ventral medial temporal cortex (AVMTC), a brain region known to be crucially involved in visual processing and object memory. In addition to their well-known function in signaling the novelty or familiarity of objects in the current environment, AVMTC neurons reflected the predictions of future novel objects, akin to the way neurons in reward-circuitry predict future rewards in order to control reward-seeking. Our data uncover a network of primate brain areas that regulate novelty-seeking. The behavioral and neural distinctions between novelty-seeking and reward-processing highlight how the brain can accomplish behavioral flexibility, providing a mechanism to explore novel objects.

## Main

Humans and other primates evolved to seek novel experiences, for example by expressing strong desires to inspect novel objects (1–7). However, the neuronal mechanisms that control novelty-seeking are poorly understood.

A dominant set of theories suggests that neurons that process reward, particularly dopamine (DA) neurons that signal prediction errors in “reward values” (8–10), also treat novelty as its own reward. This notion is mostly supported by theoretical work (11–13) which draws from the observation that when subjects are shown novel objects, functional magnetic resonance imaging detects changes in blood oxygenation in-and-around the substantia nigra (14, 15) where many DA neurons reside.

There is indeed strong evidence that DA neurons reflect the subjective value of primary, appetitive rewards (8, 16–19), and can also signal more abstract forms of reward, such as the preference for obtaining information about upcoming uncertain rewards (20) and for social interactions (21–23). However, whether or not the preference for novelty, for novelty’s own sake, is encoded in DA neurons remains unknown. Whether DA neurons, or other types of neurons, predict future novelty remains unclear because previous work studied neural response to the presentation of novel or familiar objects (24–26) but has not assessed whether and how novelty is predicted.

Theories of novelty-seeking have also arisen from the efforts of artificial-learning researchers (27), for example to construct “self-evolving” agents (2, 28–30). They propose that novelty seeking could be controlled relatively independently from reward-seeking (1,2, 28, 29). This serves to solve the famous “sparse-reward problem” by encouraging agents to seek novelty and explore it even when there are no immediate rewards to be obtained (2, 31). Consistent with this idea, a number of behavioral studies suggest that novelty-seeking and reward-seeking may be behaviorally dissociable. For example, human infants and adults and many animals exhibit novelty-seeking actions that are not related to reward expectancy (for review see (2)). To date, the question has remained whether reward-seeking and novelty-seeking are dissociable at the level of neural circuits.

Using electrophysiology, causal manipulations of neuronal activity, and detailed analyses of primate behavior, we show that the zona incerta (ZI) controls novelty-seeking of never-before-seen objects. Additional experiments showed that novelty-seeking was behaviorally and neuronally dissociable from reward-seeking. The habenula-dopamine pathway, known to reflect reward predictions and control reward-seeking, was relatively inactive during novelty-seeking, when novel objects did not predict future rewards or reward learning. This indicated that the ZI receives novelty predictions from other sources outside of the canonical circuits of motivation. High channel-count electrophysiological experiments and anatomical tracing identified the anterior ventral medial temporal cortex (AVMTC) – a brain region known to be crucially involved in visual processing and object memory (32–35) – to be a prominent source of previously unidentified novelty-seeking signals. Particularly, in addition to their well-known function in processing novelty or familiarity of objects in the current environment, AVMTC neurons carried information about predictions of future novel objects, akin to the way neurons in reward-circuitry predict future rewards in order to control reward-seeking.

This data uncovers how novelty-seeking is regulated in the primate brain through the ZI and is consistent with models that posit that reward-seeking and novelty-seeking are controlled by multiple motivational systems that are dissociable at the level of neural circuitry.

The ZI has recently gained prominence due to studies in rodents that have illuminated its role in integrating wide-ranging higher order cortical inputs (36, 37) to directly control multiple forms of motivated behavior (38–41) and mediate attentional shifting through its heterogeneous projections to the brainstem and other brain areas close to the control of action (38). In primates, the ZI has a prominent projection to the superior colliculus (42) – a key controller of gaze and attention (43, 44). However, the functional role of the ZI in primate motivated behavior has been unclear. Our data show that the primate ZI is particularly crucial for novelty-seeking, by helping to transform higher order signals about object novelty to action.

We trained monkeys to perform a *novelty seeking/inspecting behavioral task*. In *novelty-seeking* trials, monkeys could choose to shift their gaze to a familiar peripheral fractal object in order to gain the opportunity to view a novel fractal object (Figure 1A-top). To test whether neurons that predict future novel objects also respond to the onset of unpredicted novel objects themselves, the same task also included *novelty-inspecting* trials (Figure 1A-bottom). In this condition, a novel object appeared immediately at the time of fractal onset. In all trials, the amount, size, and rate of reward was not affected by the monkeys’ actions during object presentation, and the novel objects could not be used to maximize reward on subsequent trials.

**Figure 1.**
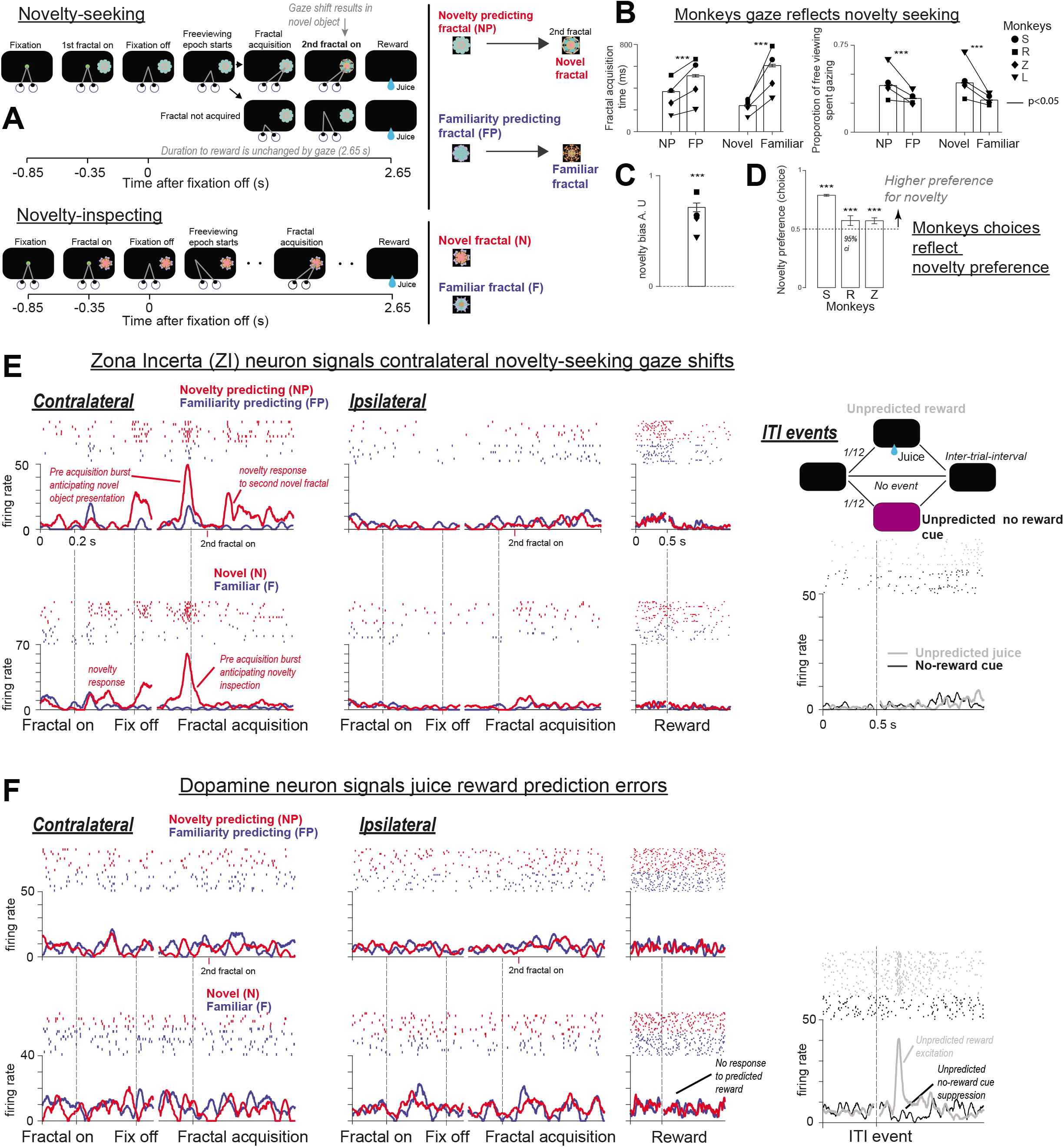
Behavior and single neurons’ activities dissociate novelty-seeking and reward-seeking. (**A**) Behavioral task diagram. (**B**) Fractal object acquisition time (**left**). Every monkey was faster to saccade to familiar objects that yielded novel objects as compared to those that yielded familiar objects (novelty-predicting objects, NP, vs familiarity-predicting objects, FP). Also, during the “free viewing” period monkeys gazed at novel objects more than familiar objects (**right**). Bars indicate mean fractal object acquisition time (**left**) and mean proportion of free viewing spent gazing at object (**right**) across all recording sessions. Symbols indicate data for each monkey separately. Error bars indicate SEM. Asterisks indicate significant differences (*** P < 0.001, Wilcoxon rank-sum test). Solid lines indicate significant differences for data within each monkey (P < 0.05, Wilcoxon rank-sum test). (**C**) Average of novelty bias index across all sessions. Each symbol indicates mean bias for each monkey. Error bar - SEM. (**D**) Monkeys’ novelty preference was assessed in novelty-choice task in which they chose among NP and FP objects. Each bar indicates each monkey’s choice rate of NP objects over FP objects. Error bars indicate 95% confidence interval. Asterisks indicate a significant deviation from chance rate (***P < 0.001, Binomial hypothesis test). **(E-F**) Task dynamics of an example ZI neuron (**E**) and dopamine neuron (**F**). Action potentials in single trials are shown by rasters, and average activity is shown by spike density functions aligned at each key trial event (**left**) and at unpredicted inter-trial events (**right**). **Left**: Activity in NP (upper, red) vs. FP trials (upper, blue) and Novel trials (bottom, red) vs. Familiar trials (bottom, blue). Trials are shown separately for contralateral and ipsilateral target presentations (relative to the recording hemisphere). Trial outcome (reward) related activity is shown combined for contralateral and ipsilateral trials. The ZI neuron (**E-left**) displayed novelty-presentation and novelty-prediction signals (in NP trials). Putative dopamine (DA) neuron (**F-left**) did not selectively respond to novelty-related events. **E-F Right**: In 1/6 of trials, unpredicted reward *or* no-reward cues occurred during inter-trial-intervals. In contrast with the ZI neuron (**E-right**), the DA neuron (**F-right**) responded with phasic activation to unpredicted reward (but not to predicted reward following task-trials; **F-left**), and it was suppressed by unpredicted no-reward cues (**F-right**). This combined pattern indicated that it signaled reward value prediction errors, but not novelty prediction errors.

In novelty-seeking trials, one of two *novelty-predicting* (NP) visual fractal objects was presented on either the left or right side of the screen. After the fixation point disappeared (‘go’ signal) the monkey was free to gaze in any manner it chose (‘free viewing’). Gazing at a NP object during free viewing caused it to be replaced by a novel object (Figure 1A-top, NP trials). Thus, the earlier the monkeys shifted their gaze to the NP object, the earlier they could gaze at the novel object. On other control trials, one of two *familiarity-predicting* (FP) objects were presented. Gazing at a FP object during free viewing caused it to be replaced by a familiar object (Figure 1A-top, FP trials). Importantly, monkeys were extensively familiarized with the NP and FP objects during training, and thus they were able to learn that NP objects were consistently associated with access to novel objects while FP objects were associated with access to familiar objects. This design allowed us to study the monkeys’ motivation to obtain novel objects by comparing how rapidly they shifted their gaze at the NP versus FP objects, in a precise analogy to conventional measures of reward seeking motivation which compare how rapidly monkeys shift their gaze to objects that deliver large or small rewards (45, 46).

Our task also included *novelty-inspecting* trials in which a novel object appeared immediately following fixation and remained on until reward was delivered (Figure 1A - bottom, N trials), and analogous control *familiarity-inspecting* trials which instead presented a familiar object (Figure 1A - bottom, F trials). All four trial types were interleaved and hence were not fully predictable. Novelty-seeking trials allowed us to study the prediction and seeking of future novel objects while novelty-inspecting trials allowed us to study neural and behavioral responses to the onset of novel objects themselves.

Previous studies showed that monkeys gaze at novel objects more than familiar objects (4, 47, 48). However, it has been mostly unclear whether or not animals are motivated by the promise of novel objects that are not yet available, an important form of novelty-seeking and prediction (49). We found that indeed in NS trials, monkeys displayed stable novelty-seeking behavior. Strikingly, all four monkeys were faster to shift their gaze onto the familiar NP objects (object-acquisition time) that yielded novel objects than onto familiar FP objects that yielded familiar objects (Figure 1B, left). Replicating previous studies, in novelty-inspecting and familiarity-inspecting trials, monkeys acquired the peripheral object faster when it was novel versus when it was familiar (Figure 1B, left) and generally gazed at novel objects more than familiar objects during “free-viewing” (Figure 1B, right; full time course of gaze is shown in Supplemental Figure 1). Hence, thus far, our task allowed us to show that monkeys predicted, and were motivated to actively seek, novel objects (Figure 1B, left). Importantly, this behavioral bias was present even though the monkeys always received the same juice reward regardless of their actions, and hence the behavioral differences across NP and FP trials could not be attributed to differences in reward expectancy. To quantify the strength of novelty-seeking behavior, we computed a novelty-bias index that quantifies the novelty related differences in object-acquisition times in trials with novel objects versus trials without novel objects. This measure isolated the influence of novelty on motivation (Figure 1C - positive values indicate positive novelty-bias). All monkeys displayed the motivation to seek novel objects (Figure 1C).

To validate that our behavioral procedure was well-suited to study whether and how neural systems signal the preference for novel objects, we verified that the response time biases (Figure 1B-C) known to reflect the level of monkeys’ motivation (45, 50) indeed match the monkeys’ preferences for novelty. To do this, we designed a behavioral procedure that let the monkeys choose between obtaining a novel or familiar object on each trial (Supplemental Figure 2). Consistent with their response time biases in NS trials, the monkeys preferred to receive novel objects (Figure 1D).

Next, using our behavioral procedure, we sought to uncover the neural mechanisms that underlie novelty seeking. To target the ZI and other brain regions, we used a combination of previously outlined electrophysiological and imaging methods (Methods, Supplemental Figure 3).

We discovered that many neurons in the zona incerta (ZI) encoded the opportunity to experience novel objects. One example ZI neuron is shown in Figure 1E. The neuron preferentially increased its activity in anticipation of gaze shifts to obtain novel objects in NP trials, and in response to novel objects themselves (Figure 1E, red trace). Also, this neuron encoded key information about the monkey’s upcoming novelty-seeking actions: (*i*) it was spatial, responding more during trials in which the peripheral object was presented on the contralateral versus ipsilateral hemi field (relative to the recording site), (*ii*) its responses were higher when its activity was aligned to the time of the fractal acquisition versus object onset, and (iii) it ramped up its activity in anticipation of the monkey obtaining the opportunity to gaze at the novel object. This neuron encoded spatial- and motor-information that is ideally suited to regulate novelty-seeking gaze shifts at visual objects.

This integration of novelty-predictions and action-control variables was strikingly similar to how basal ganglia striatal neurons integrate action signals and reward predictions to control reward seeking (51–53). This therefore begged the question. Could DA neurons send novelty predictions to the ZI?

To assess whether reward prediction error coding DA neurons signaled object novelty to the ZI, we recorded the discharge activity of putative DA neurons from substantia nigra pars compacta (SN). Neuronal identification of dopamine neurons in non-human primates followed previous studies that have been replicated in rodent SN (9, 10, 54, 55). To identify DA neurons with reward value-related activity, we augmented a small fraction of inter-trial intervals (ITIs) to include two types of stimuli that have been extensively used for this purpose in previous studies (9, 10, 54, 56–58): unpredicted rewards, and no-reward cues indicating reward omission (a brief change in screen color coupled with a sound) (Figure 1E, right). These randomly occurred during approximately 1/6 of ITIs, and never occurred within the same ITI. As in previous studies (10, 54), we expect a canonical value coding dopamine neuron to signal positive reward prediction errors following unpredicted rewards. The pattern of activity we expect is (i) excitation to unexpected rewards, (ii) relative insensitivity to predicted rewards, and (iii) inhibition to the unexpected no-reward cue.

Crucially, if a neuron that is sensitive to reward prediction errors is also sensitive to novelty-related prediction errors, it should respond to unpredicted presentations of novel objects (during NI trials), and to unpredicted presentations of objects that predict the opportunity to gaze at novel objects (during NS trials; NP). This is analogous to the precise pattern value coding DA neurons exhibit when rewards are unexpectedly delivered or omitted: responding with phasic excitation to unpredicted rewards themselves, as well as to unpredicted sensory cues that indicate reward will soon be delivered (9, 10)

An example of a putative dopamine neuron is shown in Figure 1F. This neuron showed exactly the expected, canonical response pattern to reward-related stimuli: excitation by unpredicted rewards during the ITI, no response to fully predicted rewards during the task, and inhibition by unpredicted no reward cues during the ITI. However, this neuron did not display differential activation in response to NP versus FP objects. Similarly, it was not modulated by the relatively unexpected presentation of novel objects in NI trials. Thus, remarkably, this neuron was relatively insensitive to object novelty or novelty prediction errors, despite the fact that the same neuron lawfully encoded signed reward prediction errors and it was recorded in a monkey that was motivated to seek novel objects.

We found that population level responses closely mirrored these single neurons’ results. There was prominent population of ZI neurons that was preferentially excited in contralateral NS and NI trials during the initial object presentation (Figure 2A-B; Supplemental Figures 4C). Like the example ZI neuron in Figure 1E, average activity of task sensitive ZI neurons predicted contralateral gaze shifts to NP objects and responded to the novel objects themselves in NI trials (Figure 2A; Supplemental Figure 4). The magnitude of these novelty anticipation signals in NS trials and novelty presentation related signals in NI trials were correlated across ZI neurons (Supplemental Figure 4C) suggesting that these signals are part of coherent novelty-seeking processes.

**Figure 2.**
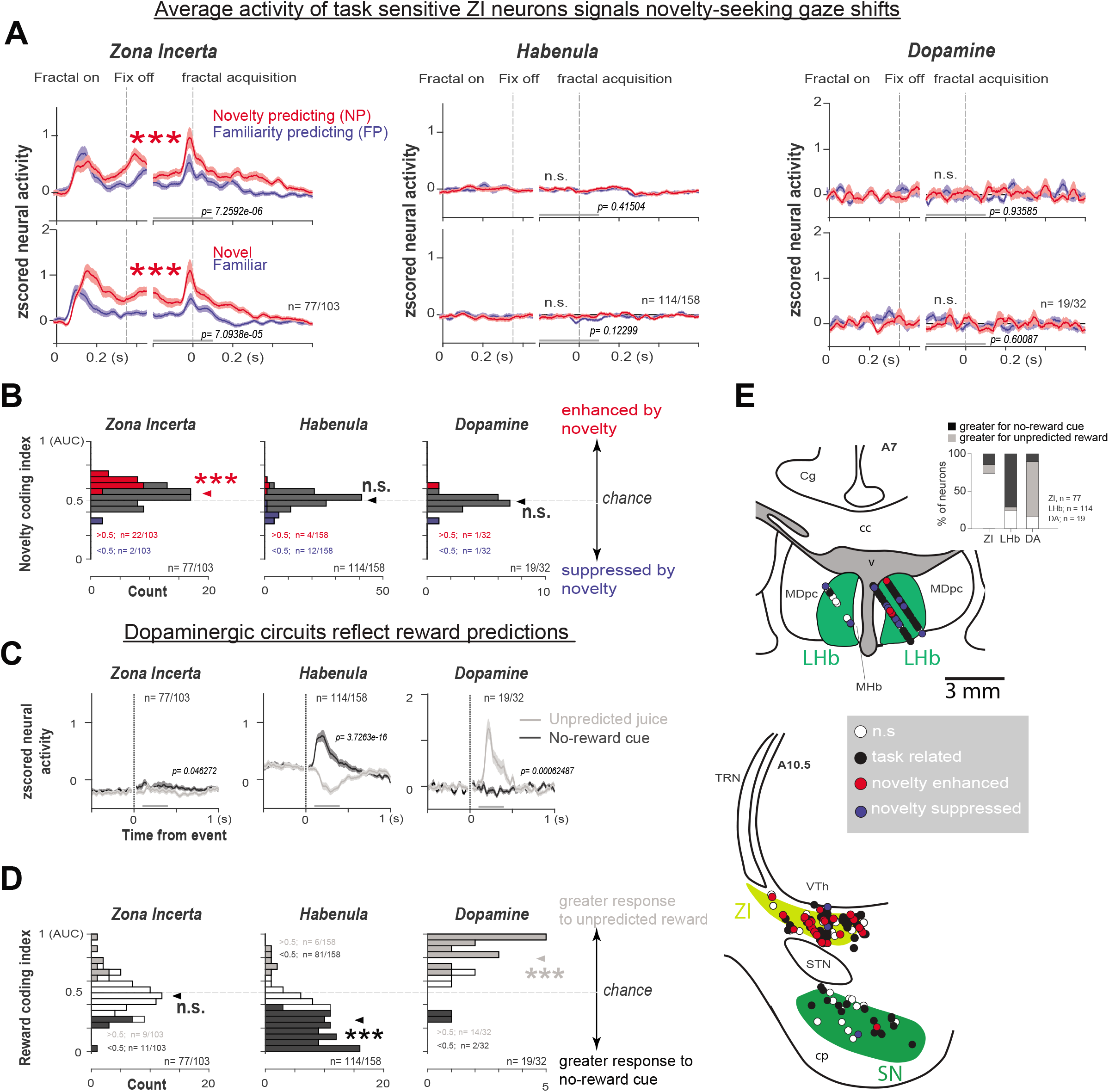
Novelty seeking is reflected by neurons in the Zona incerta (ZI). **(A)** Average activity of all recorded neurons that displayed event related variance without any further pre-selection in the ZI, habenula, and dopamine populations. Because ZI activity was spatially selective, contralateral trials are shown (spatially selectivity measures are in Supplemental Figures 4,5,7; and ipsilateral trial’s activity for ZI are shown in Supplemental Figure 4). NP (upper, red) vs. FP trials (upper, blue) and Novel (bottom, red) vs. Familiar trials (bottom, blue). Error bars denote SEM. Gray lines in each activity plot indicate window in which statistical tests were performed across average activity, and P-values of this analysis are indicated. Asterisks indicate a significant difference between responses in NP and FP trials, or Novel and Familiar trials (***P < 0.001, Wilcoxon signed-rank test). N.S. – p>0.05. (**B**) Histograms of single neurons’ novelty coding indices are shown for each area. To summarize novelty sensitivity, and increase the probability of finding significant differences in the habenula-dopamine pathway, discrimination indices (area under ROC curve; AUC) compared all novelty-related trials (NP and Novel) vs. all familiar-related trials (FP and Familiar) during the window in which habenula-dopamine neurons are known to respond to predictions and outcomes (50 milliseconds from object onset to the “go-cue”). Red and blue bars indicate neurons with a significantly larger (red) and smaller (blue) activations during novel versus familiar trials (P < 0.05, Wilcoxon rank-sum test). Arrowheads indicate mean of the distributions. Asterisks indicate significant difference from (0.5) chance (***P < 0.001, Wilcoxon signed-rank test). N.S. – p>0.05. Only the ZI (**left**) displayed a significant average novelty sensitivity (p<0.001). Also, the ZI contained more novelty-enhanced neurons than would be expected by chance (p<0.01), while the habenula (**middle**) and dopamine (**right**) did not have more novelty enhanced or suppressed neurons than would be expected by chance (p>0.05; Binomial tests assessed number of novelty enhanced or suppressed neurons among total recorded neurons). (**C**) Averaged inter-trial-interval unpredicted event activity of all neurons included in (**A**). P-values comparing activity during inter-trialinterval events are shown (Wilcoxon signed-rank test). Thick gray lines below denote analyses windows. (**D**) The ZI (**left**) contained relatively small numbers of unpredicted reward enhanced and unpredicted no-reward cue enhanced neurons; in contrast, the habenula (**middle**) had more neurons that were relatively more suppressed by reward (p<0.01) and DA (**right**) had more neurons that were relatively more enhanced by reward (p<0.01). Habenula and dopamine average discrimination (arrowhead) of reward versus no-reward events were highly significant (***P < 0.001, Wilcoxon signed-rank test), albeit in opposite directions. Across all three areas, reward and novelty discrimination indices were uncorrelated (Supplemental Figures 4, 5, 7; Spearman’s rank correlations threshold p<0.05). Indices in (**D)**: discrimination between unpredicted reward versus unpredicted no-reward events (AUCs). Reward enhanced – gray; reward suppressed – black (P < 0.05, Wilcoxon rank-sum test). (**E**) Reconstruction of recording sites. Circles indicate recorded neurons. Black filled circles - neurons with significant task-related modulations. Red and blue circles - neurons especially showing significant novelty enhanced and suppressed response, respectively (same neurons from **B**). White circles - neurons with no significant modulations. LHb neurons (recorded from A6 to A7) are shown in the upper coronal plane (A7). ZI and dopamine neurons (recorded from A9 to A12) are shown on the bottom coronal plane (A10.5). **Upper inset** indicates the proportion of neurons showing relative excitation (gray), inhibition (black), or no modulation (white) in response to unpredicted reward versus unpredicted no-reward cues. Cg, cingulate cortex; MDpc, medial dorsal thalamic nucleus, parvicellular division; LHb, lateral habenula; MHb, medial habenula; TRN, thalamic reticular nucleus; VTh, ventral thalamus; ZI, zona incerta; STN, subthalamic nucleus; SN, substantia nigra; cc, corpus callosum; v, ventricle; cp, cerebral peduncle.

Compared to the ZI, most DA neurons displayed relatively stronger responses to unexpected reward versus no-reward cues (Figure 2C-D, Supplemental Figures 5-6), but the same neurons did not show differential activity related to novelty prediction or inspection (Figure 2A-B). DA neurons showed no evidence for preferential responses to novelty-predicting stimuli or novel stimuli themselves during the initial peripheral presentation, or during free-viewing (Figure 2A; Supplemental Figures 5-6).

We further validated these results by recording in the lateral habenula (LHb) – an area exerting strong inhibitory control over numerous neuromodulators, particularly over DA value-coding neurons (59, 60). The LHb is known to respond to reward value in a mirror image to value-coding DA neurons because LHb neurons signal negative reward prediction errors (60–66) and LHb stimulation preferentially inhibits dopamine neurons that have signatures of value coding (60, 62, 67). LHb is also conclusively identifiable in monkeys based on electrophysiology (60, 62). Hence, recording in the LHb allowed us to test whether our results generalize across the dopaminergic value coding pathway. As expected, mirroring DA neurons, the majority of LHb neurons were significantly more activated by unexpected no reward cues than by unexpected reward delivery, with their average activity strongly excited by the former and inhibited by the latter (Figure 2C-D; Supplemental Figure 7). And mirroring DA neurons, the same LHb neurons were on average not significantly modulated by novelty (Figures 2A-B). Importantly, detailed neuron-by-neuron analyses revealed no relationship between novelty-related and reward-related signals in the ZI, DA, or LHb populations (Supplemental Figures 4-5 and 7). Thus, the dissociation between reward-related and novelty-related processing extended beyond DA neurons to an upstream area which plays a central role in motivational circuitry.

The locations of all neurons analyzed in Figure 2 are shown in Figure 2E. Novelty-related signals were prominently clustered in the ZI, while reward value-prediction error signals were prominently clustered in the SN and LHb.

Our results thus far show that reward-value coding dopamine neurons largely do not process object novelty in a task where it does not predict opportunities to increase reward. In contrast ZI activity strongly reflected predictions about novel objects and the actions needed to obtain them. Furthermore, these results could not simply be explained by behavioral differences across NP and FP trials such as differences in response times. To test this, we used a previously developed task in which animals are motivated to seek advance information to resolve their reward uncertainty (20, 68). This task is known to activate dopamine and habenula neurons and evokes strong behavioral preferences reflected in both choices and response times (20, 66, 68) but this task did not produce a pattern of ZI activity comparable to what we observed in the novelty seeking/inspecting task (Supplemental Figure 8). We also tested whether the bulk of our observations held true as novel objects gradually became familiar over the course of repeated exposures during a *novel-familiar learning procedure*. Indeed, in this procedure again the dopaminergic pathway was relatively insensitive to novelty, while ZI reflected the time course of novelty-to-familiarity transformations (Supplemental Figure 9).

The lack of novelty sensitivity in the LHb-DA pathway raises the question: does our task elicit novelty signals in other neuromodulator systems known to respond to novelty? Could these systems mediate novelty-seeking? We recently found that a functional class of phasically active basal forebrain (BF) neurons that is sensitive to unexpected deliveries of reward is also sensitive to the presentation of novel objects (49). We therefore tested whether these BF neurons responded to novelty in our task and carried the crucial signal for motivating novelty-seeking – a prediction of novelty in NP trials. We found that following the presentation of novel objects, BF phasic-type neurons displayed a rapid phasic activation replicating our previous work (Supplemental Figure 10A-B). But, crucially, unlike ZI, these neurons did not carry novelty predictions: it did not discriminate between novelty- and familiarity-predictive objects on novelty seeking trials. BF phasic-type neurons only displayed novelty sensitivity during NP trials well after the gaze shift and after the novel object was presented (Supplemental Figure 10C).

In sum, the neural results thus far provide evidence that the ZI uniquely encodes novelty in a manner that could mediate actions to seek future novel objects, and show that reward-seeking and novelty-seeking are dissociable at the level of neuronal circuits.

It is important to point out that our results do *not* indicate that the LHb->DA pathway is insensitive to novel objects when their novelty is a cue indicating a change in reward, or in cases in which novelty provides an opportunity for new reward-associative learning. In fact, when new objects have reward values that monkeys have not yet learned, dopamine neurons do respond to novel objects and rapidly update their value-representations as animals learn object-reward associations (18, 56, 69). In mice, unexpected novel objects that are firstly perceived as threatening rapidly activate a specific DA population in the caudal-lateral substantia nigra that are involved in processing threatening and aversive events (25). In another study, when mice were presented with neutral novel odors, responses of DA neurons were highly variable across odors and animals, in a manner that was correlated with preferences for the odors (70). And, consistent with our results, another study found no novelty-related selectivity in medial DA neurons, but did observe signals related to the subjective value of social behavior (23). These studies show that DA neurons are activated by novel stimuli in large part due to the subjects perceiving them as valuable or important (18, 25, 69, 71,72) for guiding reward- or punishment-related behaviors. Our data indicates that that LHb->DA pathway is relatively insensitive to novelty when animals are strongly motivated to seek novelty for its own sake rather than as a tool to obtain extrinsic rewards.

ZI neurons encode information that is theoretically necessary and sufficient to mediate novelty seeking gaze shifts. However, how this signal is utilized by the brain was unclear. We hypothesized that temporary disruptions of ZI circuitry would impair novelty-seeking. To test this, we injected GABAa agonist muscimol (68, 73) into the regions of ZI enriched with novelty-seeking related neurons. Based on the neuronal activity we observed in ZI (Figure 1E), we hypothesized that inactivation of this region would reduce novelty-seeking.

We compared novelty bias (Figure 1C) before and during ZI inactivations (Monkeys S and R) and found that it was indeed clearly and consistently reduced after inactivation. Importantly, the effects were highly specific and selective (Figure 3A-B) in both monkeys (Supplemental Figures 11). That is, novelty bias was quenched by the inactivation only during trials in which gaze shifts were made to the contralateral visual hemifield (relative to the injection site), but not during inter-mixed ipsilateral trials. This is important for two reasons. First, this result provides a well-established internal control (43, 68, 73) indicating that the inactivation-induced effects were spatially specific, and not due to general changes in motivation or engagement. Second, this result mirrored the strong spatial contralateral preference observed in the ZI neural activity (Figure 1E, Supplemental Figure 4B). The inactivation experiments showed that the ZI neurons that we temporarily inactivated causally impact novelty-seeking behavior. How does the ZI implement its influence on novelty seeking actions? The ZI has a powerful reciprocal relationship with the superior colliculus (42) – a key regulator of saccadic eye movements and spatial attention (44). We therefore hypothesized that the ZI regions that contain mechanisms for the regulation of novelty-seeking also have strong access to saccadic eye movements. To answer this question, we performed low intensity electrical stimulation within ZI regions that are enriched with novelty-excited neurons. Stimulation was initiated at object onset while the monkeys continued to fixate the central spot (Figure 1A) and ended 50 milliseconds before the go-cue (Figure 3C). On average, ZI stimulation facilitated upcoming contralateral object-acquisition gaze shifts (Figure 3C, right-middle), but not ipsilateral ones (Wilcoxson sign rank test; p>0.05). This was again consistent with the contralateral spatial preference in ZI neural activity and with the spatially specific effects of inactivation experiments. In contrast, stimulation of nearby regions did not facilitate saccades (Figure 3C). Off-target stimulation experiments suggested that previously developed stimulation parameters (74–78) successfully minimized current spread and off-target activation of fibers of passage (74, 78).

**Figure 3.**
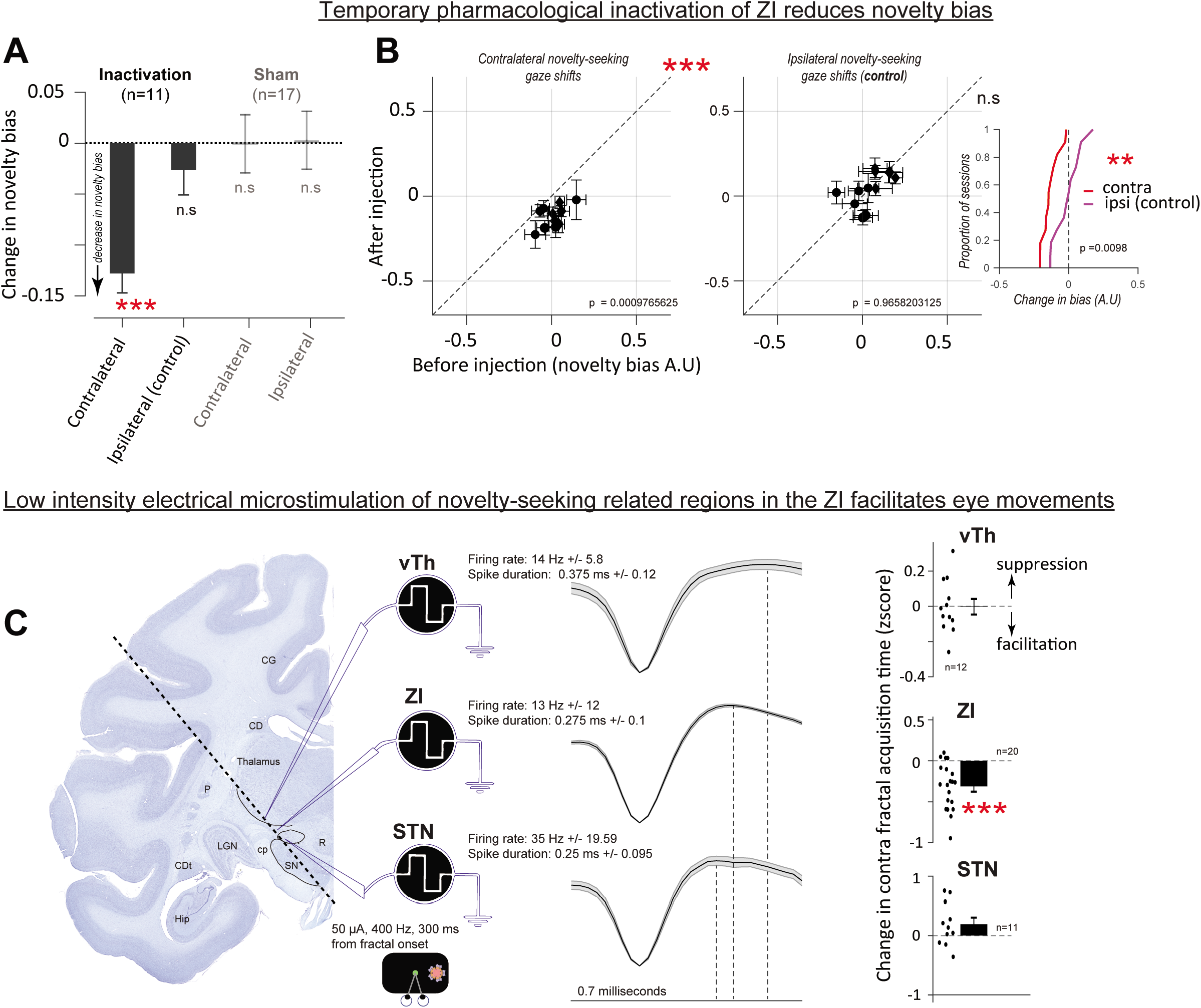
ZI is causally related to novelty seeking. (**A-B**) Temporary pharmacological inactivation of ZI regions enriched with novelty related neurons disrupts novelty seeking. (**A**) We compared novelty seeking bias (Figure 1C) before and during ZI inactivation. We found that ZI inactivation reduces novelty seeking bias only during trials in which gaze shifts were contralateral to the site of the injection (matching the spatial selectivity of ZI). Sham sessions are shown on the right for contralateral and ipsilateral trials. (**B**) Single sessions data are shown for contralateral (**left**) and ipsilateral (**right**) trials. P values comparing before and during inactivation data are indicated. Error bars denote SEM. Inset – cumulative distribution functions denote change in novelty bias for contralateral and ipsilateral trials. (**C**) Low intensity electrical stimulation of ZI regions enriched with novelty related neurons but not stimulation of neighboring brain areas facilitates contralateral target object acquisition gaze behavior. Left – example recording and electrical stimulation path shown on a Nissl stain. Middle – classical electrophysiological markers of each brain region. Average firing rates, action potential wave form shapes (+/-SEM; entire length corresponds to 0.7 milliseconds) and action potential wave form durations (dotted lines) are shown. Right – electrical stimulation of ZI (middle), but not of ventral thalamus (vTH) anatomically above the ZI or of STN anatomically below the ZI, facilitates saccadic target acquisition time (***P < 0.001, Wilcoxon signed-rank test; STN and vTH – P>0.05). Single sessions are shown as dots around the mean (bar).

In sum, our pharmacological and electrical stimulation experiments indicate that novelty-excited neurons are located in regions of the ZI that regulate novelty-seeking gaze shifts and have relatively direct access to oculomotor circuitry.

We sought to identify the neural sources of the novelty predictions that the ZI uses to anticipate and promote novelty seeking behavior. A prominent theory of ZI function is that it integrates cortical computations to directly coordinate motivation and action (38, 39, 79–82). And, our data thus far indicate that the habenula-dopamine pathway is not the source of novelty predictions in the ZI.

We therefore hypothesized that frontal and temporal cortical regions that prominently project to the ZI (37) may be the source of its novelty-predictions. Previous work in these areas has studied the responses of single neurons to the presentation of novel or familiar objects (24) but has not assessed whether and how single neurons there predict and anticipate novelty.

To identify cortical sources of novelty predictions in temporal cortex, amygdala, hippocampus, and the prefrontal cortices, we recorded thousands of neurons in 17 different brain regions (Figure 4A). This work was performed over a course of approximately four years using chronically implanted high channel count arrays with independently movable electrodes. During these neural recordings, Monkeys S and L participated in the novelty seeking/inspecting task (Figure 1A).

**Figure 4.**
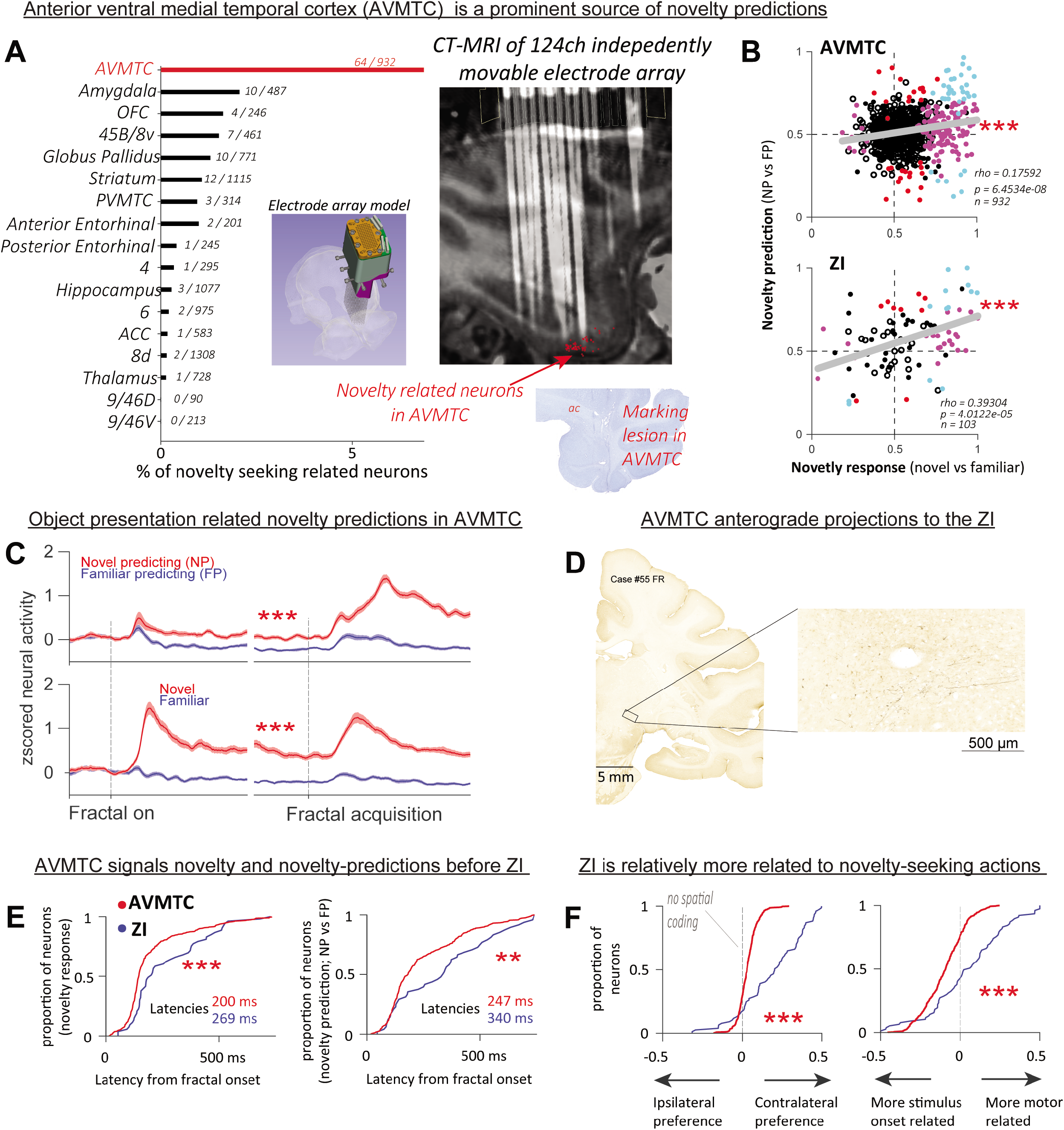
Anterior ventral medial temporal cortex (AVMTC) is a prominent cortical source of novelty prediction signals. (**A**) High channel count semi chronic array recordings revealed that AVMTC is preferentially enriched in neurons that displayed novelty-seeking control signals observed in ZI (Figure 2). Neurons that displayed task event sensitivity *and* discriminated novelty-predictions (NP vs FP) as well as discriminated novelty presentations (N vs F) with the same sign are defined as noveltyseeking neurons because they display the key signals theoretically required to drive novelty seeking (Supplemental Figure 14 shows N vs F and NP vs FP analyses separately). A neuron was defined as novelty-seeking if it passed this criterion during either contralateral or ipsilateral trials, so the results were not biased to find spatially selective (or unselective) regions. % of novelty seeking neurons is shown for 17 brain areas. The two numbers by each bar indicate the total number of neurons and the number of novelty-seeking neurons. AVMTC had higher ratio of novelty seeking neurons than each of the other areas (indicated by red bar; p<0.05; tested by 1000 permutations, Bonferroni corrected). **A - small inset on left bottom**: model of semi chronic high channel count array with 124 independently movable electrodes on monkey’s skull. **A - small inset on right bottom**: Electrolytic marking lesion in AVMTC of Monkey L at a location of a novelty-predicting related neuron. **A - large inset on right bottom**: Model of recording array is superimposed on a sagittal MRI slice of the brain. Electrodes (from a CT scan) are also shown after a single example recording session (Monkey S). Locations of AVMTC novelty-seeking related neurons across all recordings are represented by red dots. (**B**) In both AVMTC and ZI, the magnitude of novelty prediction during the NS trials was correlated with the magnitude of novelty responses in NI trials. Hence, novelty “detection” responses (in NI trials) and novelty prediction responses (in NS trials) are linked. AUC values from ROC analyses are setup such that values greater than 0.5 indicate higher discharge rates on trials with novelty predictions (y-axis) or novel object presentations (x-axis). Each dot is a neuron. Red and magenta dots indicate neurons showing significant novelty-prediction responses and novelty-presentation responses, respectively (Wilcoxon rank sum test; p<0.05). Cyan dots indicate neurons showing significance in *both*. Black dots indicate other neurons with significant task-related modulation (Kruskal-Wallis test; p<0.05) but no novelty related modulation. White dots indicate neurons with no task event modulation. Gray lines indicate least square fits. Spearman’s correlation results are reported by each scatter plot. (**C**) Average activity of AVMTC neurons pre-selected for novelty selectivity on contralateral trials during fractal acquisition epoch shown here for ipsilateral trials. Conventions are the same as Figure 2. Here, all neurons that displayed task event variance and discriminated NP vs FP trials and Novel vs Familiar trials with the same sign (either with novelty-related excitation or inhibition). (**D**) Anterograde tracer injections into AVMTC produce labeling in ZI. (**E**) **left**-latency of novelty signals in NI trials, **right**-latency of novelty predictions in NP trials. Cumulative distribution functions of neural latencies were computed from neural signals aligned on fractal object onset. Mean latencies are reported as text. ***P < 0.001, *P < 0.05, Wilcoxon rank sum test. Supplemental Figure 16B shows same analyses aligned on acquisition. The results were the same. (**F**) AVMTC detects novel and novelty-predicting objects, and ZI mediates novelty seeking actions. **F-left**– ZI neurons have relatively more spatial information than AVMTC to control action. **F-right –**consistent with latency differences, AVMTC neurons are relatively more related to the onset of visual stimuli, while ZI neurons are relatively more related to gaze motor behavior.

Across the recorded neural populations, we looked for brain regions that preferentially contained neurons that displayed the crucial pattern of activity that we observed in the ZI that is ideal to control novelty-seeking: selective prediction of the opportunity to gaze at novel objects (i.e., novelty prediction) *and* selective response to the initial presentation of novel objects themselves (Figures 1E and 2A). This screening procedure revealed that the anterior ventral medial temporal cortex (AVMTC) was preferentially enriched with novelty-seeking related neurons (Figure 4A). Similar results were also obtained from analyses of multi-unit signals (Supplemental Figure 12), and within each animal (Supplemental Figure 13).

AVMTC includes the anterior medial inferotemporal cortex and the perirhinal cortex, spanning from roughly 3mm posterior to the anterior commissure to the temporal pole (83). This region of the primate brain is known to detect novel objects when they are presented and to directly participate in object memory (32, 34, 35, 83–85). We replicated previous observations that AVMTC and many other brain regions contain single neurons that respond to the presentations of novel objects (Supplemental Figure 14). But, crucially, our data show that among the recorded areas the AVMTC in particular has a key novelty-prediction signal (Figure 4A-C; Supplemental Figure 14). As in ZI, AVMTC neurons signal the novelty of incoming sensory information *and* actively predict the opportunities to gaze at novel objects (Figures 4B-C; Supplemental Figures 15-16). Furthermore, as in Zl, the magnitude of AVMTC neurons’ novelty-prediction responses in NP trials was correlated to the magnitude of the same neurons’ activity in response to the presentation of novel objects, suggesting that the two neural signals reflect coherent processes (Figure 4B; Supplemental Figure 15).

To test whether AVMTC novelty predictions could be sent to ZI, we first tested for suitable anatomical connections. Indeed, injecting anterograde and retrograde tracers into AVMTC produced clear labeling in the ZI, indicating that the AVMTC and the ZI are monosynpatically interconnected in primates (Figure 4D and Supplemental Figure 17). This finding in primates reflects observations in rodents showing medial temporal cortical regions project to the ZI (86–88). We next tested whether AVMTC novelty-related signals occur early enough to be a source of signals in ZI. To answer this, we analyzed the latency of novelty-related signals in the activity of single neurons, during N vs F trials in which novel objects appeared at fractal onset, and the latency of novelty prediction during NP vs FP trials (Methods). Novelty presentation responses (often termed ‘novelty detection’) were earlier in AVMTC than ZI (Figure 4E-left). Similarly, novelty prediction signals were on average also earlier in AVMTC (Figure 4E-right; Supplemental Figure 16B).

This raised a key question: did AVMTC novelty predictions already contain the necessary information to guide novelty seeking actions? Or did they have to undergo sensorimotor transformations to acquire the key action-related variables present in ZI activity (such as the locations of objects and the timing of upcoming novelty-seeking gaze shifts; Figure 2 and Supplemental Figure 4)? Indeed, our analyses support this possibility. First, qualitatively, while AVMTC neurons had strong activity driven by object-presentation, ZI neurons increased their activity in anticipation of novelty seeking gaze shifts (compare Figure 4C with Figure 2A). To quantitively assess this, we calculated an index of whether activity was more strongly driven by object presentation or by the later object acquisition timing (Figure 4F-right). This approach is commonly used to assess a neuron’s or a brain region’s relative position along the sensorimotor continuum (89–91). We found that AVMTC was more visual than the ZI, displaying relatively stronger activations during object presentation than object acquisition. In contrast, the ZI was more driven by the object acquisition gaze shifts than the AVMTC (Figure 4F-right). Second, to further quantitatively assess which of the two brain areas contained more information about novelty-seeking gaze behavior, we measured single neurons’ coding of target objects’ spatial locations (Figure 4F-left). This revealed that AVMTC neurons were spatially selective, but much less so than ZI (Figure 4F-left), meaning that the directions of upcoming gaze shifts were more strongly encoded in ZI activity. Third, consistent with Figure 4F, correlations between the magnitude of neurons’ novelty-prediction responses in NP trials and the magnitude of the same neurons’ activity in response to the presentation of novel objects was present in AVMTC during both contralateral and ipsilateral trials, but only during contralateral trials in the ZI (Supplemental Figure 15).

In summary, these results show that AVMTC activity reflects predictions of novel objects and that this signal itself is relatively not action related (e.g., it contains less spatial and motor-initiation related information than the ZI). The ZI transforms novelty and novelty-prediction signals into an action control signal that contains motor parameters to seek novelty. In other words, among the AVMTC-ZI network, AVMTC identifies objects associated with novelty, and ZI transforms this signal to control behavior, likely through interactions with the AVMTC and through its strong reciprocal connections with the SC (42).

### Concluding remarks

We found that an understudied region of the primate brain, the ZI, participates in transforming novelty prediction signals into motor signals to seek novel objects. This function could be dissociated from the habenula-dopaminergic system’s role in reward processing. Particularly, LHb and DA neurons had little response to novelty in our task in which novel objects had no extrinsic value. This lack of novelty-sensitivity was observed despite the fact that animals preferred novelty, reflected in their behavioral seeking of the novelty ‘for its own sake’ (Figures 1C-D). These results show that novelty-seeking can be regulated relatively independently from reward-seeking to facilitate flexible behavioral control.

AVMTC-ZI can influence novelty seeking directly through Zl’s reciprocal connections with the superior colliculus or through its projections to the basal ganglia and prefrontal cortex (37–39, 42, 92, 93). Consistent with this idea, our data show that at the time of novelty seeking actions, neurons in motor- and sensorimotor brain areas are recruited, as would be needed to support object acquisition gaze differences across novel and familiar trial types (Figure 1 and Supplemental Figure 18).

Extrinsic value of novelty can change, from neutral to positive to negative, depending on context. And, when novelty is a tool to obtain reward, novelty signals may be reflected in the activity of medial SN and VTA value coding dopamine neurons (2, 11, 18, 25, 27, 71,94). Differentially, when novelty is a threat, it is reflected in the activity of a different group of caudal lateral SN dopamine neurons (25) that also strongly respond to other forms of external threats (25, 54). This flexible control likely relies on the fact that the brain contains distinct circuits that distinctly mediate novelty related behaviors, such as the AVMTC-ZI. Also, the interactions between novelty- and reward- seeking could take place through a cortical route, for example via prominent projections from AVMTC to the OFC (35, 95).

Our results place ZI in a unique position in the circuitry of motivated behavior in primates, and pave the way for future investigations of its other cognitive and motivational functions. The ZI receives wide input from higher-order cortical brain areas (37) and is ideally positioned to monitor and transform sensory and cognitive signals (38, 81,82) and transmit them to subcortical regions that exert powerful control over motivation (37, 92), action (37, 39, 42), and attention (38, 39). And, ZI stimulation in humans impacts motivational and cognitive states in heterogenous and complex manners (93). An important future step will be to obtain more granular information about how the distinct neuronal types (36, 37, 39) and subregions (42, 96) of the ZI cooperate to enable its functions in novelty seeking and other motivated behaviors.

Our findings expanded the role of AVMTC beyond its classical roles in processing of incoming novel objects and memory formation, and are consistent with the ideas that regions within AVMTC, process higher order object information to control subsequent processes such as association and learning (85, 97). Notably, our results show that AVMTC not only associates objects with other “known” objects, but also associates them with abstract information, such as the novelty or familiarity of future events. Thus, akin to how the reward system associates objects and actions with future rewards to control a host of reward-seeking behaviors, our data indicates that the AVMTC associates objects with future novelty to control novelty-seeking behavior, through the ZI and possibly other circuits.

## Acknowledgements

This work is supported by the National Institute of Mental Health under Award Numbers R01MH110594 and R01MH116937 to IEM, and by the McKnight Foundation award to IEM. TO, IEM, and YF performed the neuronal recordings. TO, FS, KZ, JP, analyzed the data. IEM wrote the manuscript. TO, JP, YF, and IEM discussed and revised the manuscript. AJ analyzed the anatomical data. IEM guided and conceptualized the research. We are grateful to Ms. Kim Kocher for great animal care and animal training, and to Ethan S. Bromberg-Martin and Camillo Padoa-Schioppa for giving us valuable suggestions to improve this manuscript. We are also grateful to Baldwin Goodell and Charles M. Gray for technical and scientific assistance with high channel count recording arrays.

## SUPPLEMENTAL FIGURE LEGENDS

***Figure S1. Time courses of gaze behavior in free-viewing***. Time course of proportion of free viewing spent gazing at fractal. Lines indicate averaged proportion across all sessions at each millisecond. Thickness indicates ± 1 SEM. Colors are same as in Figure 1.

***Figure S2. Novelty-choice task***. Schematic diagram of novelty-choice task.

***Figure S3. Electrode in ZI***. Coronal MRI confirming the recording location of a novelty enhanced neuron within the region of Zona incerta of monkey R. The image was acquired with a tungsten electrode (FHC) at the recording location. The electrode’s shadow is the black line (yellow arrow). CD – caudate nucleus; CG – cingulate; vTH – ventral thalamus; ZI – Zona incerta.

***Figure S4. Supplemental analyses on ZI neurons***. (**A**) Average ZI activity in response to trial events. All conventions are as in Figure 2. Time windows for analyses of average activity across neurons are denoted by gray bars. (**B**) Histogram of spatial selectivity of individual ZI neurons. The selectivity was quantified as the AUC comparing fractal acquisition related activity during contralateral vs. ipsilateral trials. Light gray bars indicate neurons with a significant difference across the two conditions (P < 0.05 Wilcoxon rank-sum test). Arrowhead indicates mean of the distribution. Asterisks indicate the distribution was significantly different from chance (***P < 0.001, Wilcoxon signed-rank test). Numbers of cells preferring contralateral fractal trials (>0.5) and preferring ipsilateral fractal trials (<0.5) are indicated. (**C**) Scatter plots show the relationship between the magnitude of neural novelty predictions (NP vs FP; y-axis) and neural novel object presentation related responses ‘novelty detection’ (Novel vs Familiar; x-axis). Data are shown for contralateral and ipsilateral trials separately. AUC values from ROC analyses are setup such that values greater than 0.5 indicate higher discharge rates on trials with novelty predictions (y-axis) or novel object presentations (x-axis). Similarly, values less than 0.5 indicate higher discharge rates for trials with familiar objects. All conventions are the same as in Figure 4B. As would be expected from strong spatial tuning (**A-B**), ZI neurons display a clear relationship between novelty prediction and novelty detection related responses on the contralateral side. These indicate that in ZI, novelty selectivity and novelty prediction are linked during contralateral trials. To obtain accurate fractal onset related novelty selectivity for each neuron before the monkeys made saccades, we narrowed the window for calculating these indices, restricting the comparisons to the time window before the go cue. (**D**) Magnitude of novelty-related responses (NP+Novel vs FP+Familiar; x-axis) were not related to the magnitude of neural discrimination of unpredicted ITI events.

***Figure S5. Supplemental analyses of dopamine neurons***. (**A**) – Average lacks spatial selectivity. (**B**) – Novelty task related activity lacks coherent structure. (**C-left**) – excitatory DA responses are driven by events with a positive reward value. And, the magnitude of positive value coding across distinct trial events is correlated (**C-right**). Particularly, because a trial start cue indicates the timing and opportunity to gain reward, dopamine activity displays trial start cue excitation (x-axis) (67, 98). The magnitude of this excitation is correlated to ITI event positive value coding (y-axis) on a neuron-by-neuron basis indicating that on average value coding in the dopamine population reflects coherent positive value coding process across many diverse task events (67, 99). (**D**) Consistent with a lack of coherent novelty selectivity (**Figure 2**), magnitude of novelty signals was not correlated to value related events. Like in other monkey studies (72, 100, 101), dopamine neurons in our task were not spatially selective (**A**). So, throughout this figure contralateral and ipsilateral trials are analyzed together. Conventions are as in Figure S4.

***Figure S6. Additional data analyses on dopamine neurons, related to Figure S5***. (**A**) Task event dynamics of no-reward “sensory-cue” suppressed (**A**) and no-reward “sensory-cue” enhanced dopamine neurons (**B**). The two averages were of neurons with ROC values in right-bottom and right-upper quadrants, respectively, of Figure S5C, left. (**C**) Comparison of mean waveforms (left) and cumulative distributions of discharge rates (right) between both types. Gray areas in waveform indicate ± 1 SEM. N.S. - indicates no significant differences between the two groups (P>0.05, Wilcoxon ranksum test). (**D**) Reconstruction of recording sites. Circles indicate locations of recorded putative DA neurons. Gray and black indicate no-reward sensory cue suppressed DA neurons and no-reward sensory-cue enhanced types, respectively. White – DA neurons with task event related activity.

***Figure S7. Supplemental analyses of lateral habenula neurons***. (**A**) – average lacks spatial selectivity. (**B**) – Novelty task related activity lacks coherent structure. (**C-left**) – LHb excitatory responses are strongly driven by the unpredicted no-reward cue, an event with negative reward value. And, the magnitude of negative value coding across distinct trial events is correlated (**C-right**). Particularly, because a trial start cue indicates the timing and opportunity to gain reward, habenula activity displays trial start cue suppression (67, 98). The magnitude of this suppression is correlated to ITI event negative value coding on a neuron-by-neuron basis indicating that negative value coding in the habenula reflects coherent negative value coding across many diverse task events (67, 99). (**D**) Consistent with a lack of coherent novelty selectivity (**A**), magnitude of novelty signals was not correlated to negative value related events. Like in other monkey studies (60, 63), habenula neurons in our task were not spatially selective (**A**). So, throughout this figure contralateral and ipsilateral trials are analyzed together. Conventions are as in Figure S4.

***Figure S8. ZI activity in reward uncertainty motivated-behavior***. ZI activity in a task that measures the motivation to reduce reward uncertainty (“info seeking task” (20, 66, 68, 102)). (**A**) Task closely resembles previous work and has the same time course as novelty-seeking task (Figure 1A). Here, there are three trial types, info uncertain trials (red), no info uncertain trials (blue), and safe trials (black). In info trials (1/3 of trials), an info predictive fractal, which predicted the future appearance of one of two *informative* fractals, appeared at fractal onset. As soon as the monkeys gazed at this peripheral fractal, it was immediately replaced by one of two other informative fractals. One informative fractal predicted a big amount of juice reward whereas another informative fractal predicted a small amount of juice reward. Hence, these secondary fractals fully informed the animals about the value of the future outcome. On contrary, in no-info trials (1/3 of trials), another fractal predicted the presentation of one of two non-informative fractals. If the animals gazed at it, the fractal was immediately replaced by one of two non-informative fractals. Irrespective of which non-informative fractals appeared, big or small juice rewards were delivered at the time of the outcome with 50% chance. So, these non-informative fractals never informed the monkeys about the future. Finally, the remaining 1/3 of trials were safe trials in which the same amount of reward was delivered. The two secondary fractals did not predict different reward amounts. Importantly, the expected values (EV) of all the trial types were fixed. The outcomes were delivered at the same time, irrespective of monkeys’ gaze behavior. We previously found that monkeys were highly motivated to resolve their reward uncertainty, displaying shortest fractal acquisition time during Info trials (68). We replicate this here (**B**). Data is shown combined across Monkey S and R, but both monkeys displayed the shortest Info fractal acquisition times (p<0.05). Asterisks indicate significant differences (P*** < 0.001, Wilcoxon rank-sum test). Monkeys response times indicated a strong subjective preference to receive uncertainty-reducing information during Info trials. (**C**). Averaged contralateral trials’ activities of ZI neurons (left, info-seeking task; right, novelty-seeking task; Figure 1). Activities are aligned on fractal acquisition. In info seeking, red, blue and black lines indicate info, no-info and safe trials, respectively. Right - red and blue lines indicate combined novel (NP + novel) and combined familiar (FP + familiar) trials, respectively. Error bars represent SEM. To study the precise differences in the time course and ZI dynamics we used a 5 millisecond Gaussian kernel to convolve the spiking activity. There are several key takeaways from the results. First, the pre-fractal acquisition activity during info-seeking trials did not reflect response time biases of the monkeys. Particularly, there was no significant difference between info and no-info trials, which showed the greatest relative differences in response time biases (**B**). This result indicated that the ZI neurons we recorded do not drive the info-seeking response-time bias in this task, and more importantly, do not simply reflect any difference in response time distributions. Instead the information seeking bias in this task was previously found to be causally related to a cingulate-basal ganglia circuit (68) and to the phasic activations of the dopaminergic system that signals behavioral preferences for the opportunity to resolve uncertainty (20, 66, 102). In novelty seeking trials, we see clear differentiation in ZI before the action that reflects the monkeys’ behavioral biases (Figure 1). Following fractal acquisition, as the monkeys awaited to receive information, ZI activity showed differential activation for Info and No info trials which we previously identified as a signature of information anticipation (31,68, 102). In summary, ZI activity differences in novelty-seeking task are not simply a reflection of response time differences.

***Figure S9. Behavioral and neural modulations across novelty-familiarity transformations***. (**A**) task diagram. (**B**) Monkeys expressed classical behavioral effects reflecting the novelty-familiarity transformations of objects. Duration spent gazing at the objects during free viewing decreased as a function of object presentation number. Trials are shown in bins of object presentation number (x-axis). (**C**) Same convention as **B**. Neural activity is shown for distinct neural groups. Dopamine and habenula did not differentiate novel and familiar objects across novelty-familiarity transformations. On average, however, the ZI reflected changes in the animals’ behavioral familiarity in **B**. In B-C, bins without an asterisk were not significantly different across novel and familiar trials (p>0.05).

***Figure S10. Basal forebrain activity does not predict novel objects***. (**A**) Same conventions as in Figure S4. Consistent with our previous report, basal forebrain phasic activated neurons (BF) (49) responded to the presentation of novel objects in novelty-inspecting trials (bottom) and following NP fractal acquisition in novelty seeking trials (top). However, unlike ZI, these BF neurons did not show any anticipatory activity in NP and FP trials before the novel object was presented. These data are consistent with our previous proposal that BF signals external motivationally salient events (49). (**B**) Neuron-by-neuron histogram indicates that most phasically active BF neurons were excited by the presentation of novel objects (**B-left**) and not spatially selective (**B-right**). (**C**) Novelty responses in novelty inspecting trials (x-axis) were not correlated with discrimination of NP versus FP objects. Most neurons did not discriminate NP and FP objects. BF phasic neurons were identified as in previous study (49). BF neurons in our task were not spatially selective (**A**) (103). So, throughout this figure contralateral and ipsilateral trials are analyzed together. Conventions are the same as in Figure S4.

***Figure S11. Cumulative distributions of ZI inactivation related biases on a session by session basis***. Same conventions as Figure 3.

***Figure S12. Multiunit responses across brain areas***. % of multi-unit sites with novelty-seeking signals. The key result of Figure 4A is replicated. The two numbers by each bar indicate the total number of sites and the number of sites with novelty-seeking signals. Conventions and definitions of novelty-seeking signals are the same as in Figure 4A.

***Figure S13. Percentage of neurons displaying novelty-seeking related signals (same as Figure 4A) for Monkey L and S***. Conventions and definitions of novelty-seeking signals are the same as in Figure 4A.

***Figure S14. Percentage of neurons displaying novelty presentation responses and novelty predictions shown separately across brain areas***. Previous work studied novelty responses in single neurons by presenting animals novel and familiar objects and measuring differential neural responses to them. Here, we performed this analysis on an area by area basis (% of neurons shown in blue) during novelty-inspecting trials (Figure 1A). As in previous studies, many brain areas show significant novelty presentation related responses (Red and green asterisks denote p<0.01 and p<0.05 thresholds respectively; Binomial test). On the other hand, our task also measured novelty predictive activity – that is activity related to the anticipation of novel objects in NP versus. FP trials (% of neurons shown in red). Among the recorded regions, only AVMTC displayed more neurons that displayed novelty prediction than would be expected by chance. Analyses were done such that chance level is conventional 5% threshold. For each cell we asked if a neuron discriminated two conditions (N vs F for novelty presentation test; or NP vs FP for novelty prediction test) during target onset epoch. If a neuron displayed selective activation (p<0.025) on either *contralateral* or *ipsilateral* trials we counted it as significant. Therefore, after correcting for multiple comparisons (0.025 x 2) the chance level was 5%.

***Figure S15. Correlations of novelty-seeking and inspecting signals*** in AVMTC and ZI during task events. Contralateral (**A**) and Ipsilateral (**B**) trials are shown separately aligned on fractal onset and fractal acquisition. Same conventions as in Figure 4B. AVMTC displays correlations during either contralateral or ipsilateral trials, but ZI does so only during contralateral trials.

***Figure S16. Activity of AVMTC neurons***. (**A**) Average z-scored activity of cells in Figure 4C but shown for contralateral and ipsilateral trials. (**B**) Same convention as Figure 4E. Analyses were performed on activity aligned on target acquisition here rather than target onset.

***Figure S17. Retrograde and anterograde injections into AVMTC and other temporal regions produce labeling in ZI***. Consistent with rodent literature (86–88), retrograde injection into perirhinal cortex and amygdala produces labeling in the ZI (upper left). Similarly, anterior perirhinal / ventral temporal pole anterograde tracer injection produces results in ZI labeling (upper right). For each case, injection site and label are shown.

***Figure S18. Gaze action related regions are recruited with AVMTC-ZI to support novelty seeking***. Ultimately to guide action gaze control centers in the brain must be recruited - to aim and move the eye to novelty-predicting targets. Therefore, we hypothesized that dorsal prefrontal cortex and the basal ganglia could be recruited later in the trial as the monkey shifts gaze to the novelty-predicting target fractal. To test this we repeated (**A**) same analysis as in Figure 4A triggered on fractal acquisition. This analyses would therefore also detect neurons whose activity are correlated with response time biases during NvsF trials and during NP vs FP trials. In other words it would not be selective for novelty-related neurons per se. As in Figure 4A, here, AVMTC is most enriched in novelty seeking neurons, but additional regions are recruited during fractal acquisition actions: 45b/8v and globus pallidus (GP). These regions are known to be related to spatial gaze control and spatial attention (91, 104–106) and their activity is directly to saccadic behavior. To further study how the basal ganglia is recruited during novelty seeking, we also analyzed neurons in subthalamic nucleus and substantia nigra pars reticulata (SNr) – two additional regions implicated in the regulation of gaze (51, 107, 108) that we use to localize DA and ZI neurons. We found that unlike ZI and AVMTC, during fractal onset, these 4 regions do not contain prominent groups of novelty prediction neurons (**B-left, and see Figure S14**). And, only GP shows a very weak correlation between novelty inspecting and predicting signals. In contrast, during fractal acquisition (**B-right**) 45B/8v, GP, and SNr display significant correlations between novelty inspecting and predicting signals suggesting that they may become involved in novelty-seeking particularly during the execution of novelty-seeking gaze behaviors.

***Figure S19. Recording locations relative to the anterior commissure (AC). (A)*** cell # : number of cells in the region. mean(L) - AC(L) : mean of lateral coordinates of the cells in the region, referenced to AC. median(L) - AC(L) : median of lateral coordinates of the cells in the region, referenced to AC. std(L) : standard deviation of lateral coordinates of the cells in the region. |mean(L) - min(L)| : distance of most medial site in the region to mean. |mean(L) - max(L)|: distance of most lateral site in the region to mean. mean(A) - AC(A) : mean of anterior coordinates of the cells in the region, referenced to AC. median(A) - AC(A) : median of anterior coordinates of the cells in the region, referenced to AC. std(A) : standard deviation of anterior coordinates of the cells in the region |mean(A) - min(A)|: distance of most posterior site in the region to mean. |mean(A) - max(A)|: distance of most anterior site in the region to mean. mean(D) - AC(D) : mean of dorsal coordinates of the cells in the region, referenced to AC. median(D) - AC(D) : median of dorsal coordinates of the cells in the region, referenced to AC. std(D) : standard deviation of dorsal coordinates of the cells in the region |mean(D) - min(D)|: distance of most ventral site in the region to mean. |mean(D) - max(D)|: distance of most dorsal site in the region to mean. **(B)** Same as **A** but instead of information about all neurons, only information about novelty-seeking related neurons is indicated aligned on fractal onset (top) and fractal acquisition (bottom). Note that striatum is shown as a 3D plot in Figure S20 because this table cannot give the true sense for its unique shape and the location of neurons within it. The striatum wraps around from the frontal to the temporal cortex along the anterior-posterior axis (Figure S20).

***Figure S20. Posterior ventral striatum is enriched with neurons that respond to novel objects, but not preferentially in novelty seeking neurons***. Two views of all striatal neurons are shown relative to the AC (anterior commissure) along the dorsal-ventral, medial-lateral, and anterior-posterior axes. Filled color definitions are the same as in Figure S15. Because this plot does not show discrimination values (AUC) but indicates location, to be complete, we also show neurons that discriminated N vs F and NP vs FP in different signs in blue. Dotted line represents AC along the dorsa-ventral axis. Statistical tests were done on activity aligned on fractal acquisition to match Figure S18. Novelty responding neurons (magenta) are mostly found in posterior ventral regions of the striatum. This replicates the work of Yamamoto, Monosov, Yasuda, and Hikosaka (2012, Journal of Neuroscience). There were 115/731 novelty responding neurons (magenta; below the AC) in the posterior ventral regions, while above the AC, there were only 33/384 (regions below AC were significantly more enriched with novelty responding neurons; p<0.001). Novelty-seeking related neurons (cyan) and neurons that only discriminated NP vs FP (but not novel versus familiar; red) were relatively similarly uncommon in dorsal and posterior ventral striatum (Figure 4A and Figure S18). There were 7/384 novelty-seeking related neurons in the dorsal regions and 9/731 of the posterior ventral regions; and there were 5/384 cells that discriminated NP vs FP (but not novel versus familiar objects) in the dorsal regions and 30/731 in the posterior ventral regions. Overall, Figures S18 and S20 suggest that the AVMTC-ZI system can interact with several basal ganglia and prefrontal circuits to mediate novelty seeking in a task in which novelty has no extrinsic reward value.

## METHODS

### General procedures

Adult male rhesus monkeys (*Macaca mulatta*) were used for the electrophysiology experiments. All procedures conformed to the Guide for the Care and Use of Laboratory Animals and were approved by the Washington University Institutional Animal Care and Use Committee. A plastic head holder and plastic recording chamber were fixed to the skull under general anesthesia and sterile surgical conditions. For Monkeys R and Z, large neuronal recording chambers were tilted and aimed at SN, LHb, and ZI. Their anterior-posterior extent included other regions of interest such as the BF. After the monkeys recovered from surgery, they participated in the behavioral and neurophysiological experiments.

For acute recording expeirments, recording sites were determined with 1 mm-spacing grid system and with the aid of MR images (3T). This MRI-based estimation of neuron recording locations was aided by custom-built software (PyElectrode, (109)) and histology. Single-unit recording was performed using glass-coated electrodes (Alpha Omega), epoxy-coated electrodes (FHC), and 16 and 32 channel linear arrays (v-probes, Plexon). In particular, for ZI and dopamine neurons, we used custom-modified epoxy electrodes with 1.2 to 2.5 MΩ impedance (FHC). Electrodes or linear arrays were inserted into the brain through a stainless-steel guide tube and advanced by an oil-driven micromanipulator (MO-97A, Narishige). Signal acquisition (including amplification and filtering) was performed using Plexon 40kHz recording system. In these experiments, action potential waveforms were identified online by multiple time-amplitude windows, and the isolation was refined using offline clustering on the first three principal components and a measure of non-linear energy (Plexon Offline Sorter).

For Monkeys L and S, we implanted semi-chronic high channel count recording drives (LS124; Gray Matter). To aim the micro drives, we first acquired 3T magnetic resonance images of the monkeys’ brain. We used these MRIs to aim the two micro drives towards the regions of interest, including the prefrontal cortex and the temporal cortex. We then attached MRI compatible chambers to the skull, as usual using MRI compatible ceramic screws (Thomas). After the animals recovered, we performed MRI with fiducials such that we could estimate and reconstruct the path of each electrode (68, 103, 109–111). Following this pre-operative confirmation, we implanted both animals with 124-channel micro drives. These are detailed here: https://www.graymatter-research.com/documentation-manuals. Following craniotomy, we sealed the chamber and used a port to assess whether bacterial growth occurred. Following this safety precaution, we implanted the recording drives containing the electrodes and lowered all channels immediately beyond the dura. In this way, we minimized the impact of post-op dura thickening on the electrode impedance and trajectory. Data from electrode-channels were included in the study if (1) post-op CT images showed that the electrodes were in the brain and were following a trajectory that could be reconstructed, (2) if the electrode-channel produced single units during the history of the array neuronal recordings, and (3) if the post-op impedance was >0.2MΩ or single units were observed. This approach produced 108/124 channels in Monkey L and 124/124 channels in Monkey S. A key difference in success was due to the use of glass coated electrodes (Alpha Omega) in Monkey S versus thinner epoxy electrodes in Monkey L (FHC). Recording locations, including particularly the locations of novelty-selective neurons, were verified in several ways. First, we placed electrolytic marking lesions in Monkey L (74, 112, 113). Second, for both animals, we acquired sequential CT images. These images were registered to the MRI and the locations of the electrodes were directly visualized. Third, as we moved we used functional and anatomical landmarks classically employed by our and other laboratories (e.g. electrophysiological patterns of brain areas like the globus pallidus, striatum, anterior commissure, ventricles, lateral geniculate nucleus) to further verify electrode locations. Hence, like other recent studies using similar technology, for reconstruction, we combined histology, imaging, classical electrophysiological methods (110, 111, 114). The brain regions in the MRI were defined exactly as in previous studies (110, 111, 114, 115) with two exceptions. Posterior 45B and ventral 8 (8v) are hard to differentiate and their separation are contentious. We therefore grouped them together in this study (Supplemental Figure 19). AVMTC included perirhinal cortex and medial TE (Figure 4A) and was defined widely along the anterior-posterior axis from ~ −3 mm relative to the anterior commissure, spanning to the temporal pole as in previous studies (Supplemental Figure 19). Previous functional studies have grouped these regions because MRI and histology do not agree on how to differentiate their precise borders along the anterior-posterior axis. The actual locations of novelty-seeking neurons are shown in Figure 4A.

The semi chronic drive contained electrodes with 1.5mm spacing. Signal acquisition (including amplification and filtering) was performed using Plexon 40kHz recording system. Action potentials were identified in two manners. First, we used offline manual sorting (Plexon Offline Sorter). Second, we deployed a semi-supervised template matching based algorithm (Kilosort2) to sort the data and then corrected the results further to avoid over-splitting. To verify the key results of our approach, we also analyzed multi-unit activity (MUA) – a bulk, average, unbiased measure. We defined MUA as signals that passed -2 SD but did not cross -3 SD.

To identify substantia nigra (SN), we used standard landmarks such as subthalamic nucleus, ZI, and the thalamus. We identified putative DA neurons in SN based on classical electrophysiological criteria across monkey studies that have been replicated in optogenetically identified SN DA neurons (9, 10, 54, 55): 1) a low background firing rate at around 5 spikes/s, 2) a wide spike waveform in clear contrast to neighboring neurons with a high background firing rate in the substantia nigra pars reticulata, and 3) a phasic excitatory activity caused by an unexpected reward delivery or trial start cue (116). Because medial DA populations in the VTA are harder to identify online by these criteria (117), we concentrated on the SN. We also recorded in the LHb, which is known to reliably mirror relatively medial motivational value coding DA neurons’ activities (60, 62).

### Histology

After the end of some recording sessions, we made electrolytic microlesions at the recording sites in Monkey L (~20 μA and 30 s). The monkey was deeply anesthetized using sodium pentobarbital and perfused with 10% formaldehyde. The brain was blocked and equilibrated with 10% sucrose. Frozen sections were cut every 50 μm in the coronal plane. The sections were stained with cresyl-violet. Monkeys that additional participated in tracer injections were perfused in a similar way.

### Tracer related histology

Two male macaque monkeys were used. These antero-retrograde cases come from the collection of anatomical tracers from the laboratories of Dr. Joel Price that have been digitized by the Monosov laboratory for further analyses. For those injections, Fluoro Ruby (case # 55; 1 μl @ 10%) and Lucifer Yellow (case #58; 0.5 μl @ 10%) were used. Case #55 was published in a previous work by Kondo et. (2003). With the permission of Dr. Price, we re-examined and analyzed the case in relation to the anatomical connections between ZI and anterior ventral medial temporal cortex (AVMTC). The results from case #55 are reported here for the first time. For surgeries and MRI scan anesthesia was induced by ketamine injection (10 mg/kg) and maintained with a gaseous mixture of oxygen, nitrous oxide, and halothane. Animals were given analgesic (buprenorphine, 0.1 mg/kg, i.m.) post-surgery. Craniotomies were performed at the stereotaxic coordinates of the injection sites. Prior to the injections, recordings were performed along the trajectory of the injection site. This procedure helped refine the position of the injection site by controlling for the grey and white matter, sulci, as well as the bottom of the brain. 1 μl of Fluoro Ruby (FR; @ 10%) and 0.5 μl of Lucifer Yellow (LY, @ 10 %) were injected to the AVMTC of case #55 and case #58 respectively. Tracers were injected through micropipettes using air pressure. After each injection, the pipette was left in place for 30 minutes. After 14 days following the surgery, the animals were deeply anesthetized (ketamine, 10 mg/kg), overdosed with sodium pentobarbital (25–30 mg/kg), and perfused with phosphate-buffered saline followed by 4% paraformaldehyde solutions, first at pH 6.5, then at pH 9.5, and finally at pH 9.5 with 10% sucrose. The brain was removed and transferred through 10, 20, and 30% sucrose solutions in phosphate buffer at 4°C. After, Brains were frozen in isopentane and dry ice and later cut into several series (12 and 10 series for case #55 and case #58 respectively) of coronal sections at 50 μm thickness. Both FR and LY were processed immunohistochemically with an avidin-biotin-horseradish peroxidase technique.

### Task

The behavioral task is displayed in Figure 1A. The task began with the appearance of a small orange circular trial start cue at the center of the screen. The monkeys had to fixation this spot for 0.5 seconds. If the animal did not do this within 5 seconds, the trial aborted and the inter trial interval (ITI) started. Following successful fixation, a peripheral visual fractal object stimulus appeared 10 degrees visual angle from the fixation spot. The monkey had to continue to fixate the central fixation spot for 0.35 seconds, or 0.5 seconds. Then, the central fixation spot disappeared and the monkeys were free to gaze anywhere they wished because reward would always be delivered. 50% of the trials were termed novelty-seeking trials (Figure 1A, upper). During one half of these trials, one of two familiar objects appeared. These objects predicted the delivery of a novel object contingent on the animal behavior. That is, if the animals gazed at these peripheral familiar visual objects, they were immediately replaced by a novel object which remained on the screen until the outcome. We termed these novelty-predicting object trials (NP). During the other half of the novelty-seeking trials, one of two familiar objects was presented, and if the animal gazed at them, instead of a novel object, one of two familiar objects was shown. We termed these familiarity-predicting object trials (FP). The task-timing and reward parameters of these two trial types were precisely the same. Another group of trials we termed novelty-inspecting trials (Figure 1A, bottom). These constituted the other 50% of the trials. During these trials, the peripheral objects presented following trial start fixation were either novel (on half of the novelty-inspecting trials) or one of two familiar objects. The novelty-seeking and inspecting trials were not blocked. So, in sum during novelty inspecting and seeking trials, following the trial start fixation epoch, the animals experienced a novel object with 25% chance and a familiar object that could deliver a novel object (NP trials) with 25% chance.

A distinct group of trials began with a pink fixation spot. During these trials the monkeys experienced either a novel object (50%) or one of two familiar objects. However, the novel object was not “regenerated” during the experimental session. That is, it underwent novelty-familiarity transformation because the novel object remained the same and re-appeared throughout the recording session. These trials were used to gain additional evidence that novelty directly mediates gaze behavior. They also further tested whether the dopaminergic system differentiates novelty-familiarity. The timing and reward statistics of these trials were the same as the other trials described above.

In one 6th of trials, unpredicted rewards or unpredicted sensory events, termed unpredicted no-reward cues, were delivered during the ITI (ranging from 0.7s to 1.5s from ITI start). During a single ITI with an unpredicted event, the monkeys experienced only one type of ITI event (reward *or* no-reward cue).

We further verified monkeys’ novelty preferences in a distinct choice task (Supplemental Figure 2). Here, the animals chose between novelty-predicting and familiarity-predicting objects (same as in Figure 1A in novelty-seeking trials). In this way the monkeys could choose to obtain a novel or familiar object. Reward was not dependent on whether the monkeys chose to receive familiar or novel objects. Hence, the task measured the monkeys’ preference to obtain the opportunity to gaze at a novel object, not their preference for rewards.

### Statistical analyses of neuronal activity

All statistical tests are specified wherever p-values are reported and were non-parametric unless otherwise noted. When permutations were used, the number is specified. Significant task responsiveness was defined as variance in neural activity across all task events, including ITI events. To determine whether recorded neurons had significant task event related modulations, we computed p values by comparing activities within the time windows 50 to 350 ms from object onset, -200 to 100 ms from gaze object acquisition, and 100 to 400 ms from ITI events (Kruskal-Wallis test; p<0.05). The windows were chosen such that they included key neuronal modulations in each area and were wide enough to avoid bias towards a particular response pattern (e.g. phasic versus tonic). The comparisons were done across ipsilateral and contralateral NP, FP, Novel, and familiar trials; and in the ITI window across unpredicted reward, unpredicted no-reward cue, and no-event activity. The p-values were then combined (120, 121). This non-parametric method has less assumptions then parametric methods. Nonetheless, we also cross-validated it by a parametric general linear model approach which yielded similar key results for each brain area.

Neural activity was converted to normalized activity as follows. Each neuron’s spiking activity was smoothed with a Gaussian kernel (mean = 50 ms, or 5ms for temporal analyses in Supplemental Figure 8) and then z-scored. To z-score the activity, the neuron’s average activity time course aligned at post fixation peripheral object onset was calculated for each condition. These average activity time courses from the different conditions were all concatenated into a single vector, and its mean and standard deviation were calculated and used to z-score the data. Henceforth, all future analyses converted that neuron’s firing rates to normalized activity by (1) subtracting the mean of that vector, (2) dividing by the standard deviation of that vector (68). To quantifying the novelty-motivated object-acquisition response time bias (novelty bias), we created an index that was computed from the differences in object-acquisition times in NS and NI trials:

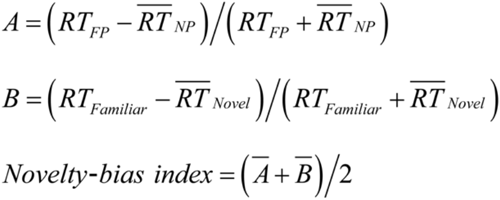

Where RT_NP_, RT_FP_, RT_Novel_, RT_Famliliar_ indicate object-acquisition times in NP, FP, Novel and Familiar trials, respectively. The index was computed separately for each session.

For semi chronic array recording, to avoid the silencing of trends by statistical thresholds, we report the raw cell count numbers, verify the results with MUA analyses, and assess whether the relative strength of novelty-seeking related signals across brain areas identifies the same brain areas as prominently carrying novelty-seeking related signals. All of these converge on AVMTC. Because in ZI we found that novelty signals were correlated in novelty-seeking and inspecting trials, we defined novelty seeking neurons as those task-related cells that displayed novelty preference in both trial-types (rank sum test; p<0.05) with the same coding sign (e.g. inhibited by novelty in both or excited by novelty in both). A neuron could be classified as novelty-seeking related if it passed our criteria on either contralateral or ipsilateral trials to the recording site. This classification identified AVMTC as a prominent source of novelty-seeking signals (Figure 4A).

Across all experiments, to derive neural selectivity indices that measured the strength of discrimination among task events, we used ROC area (area under curve, AUC) to distinguish spike counts across groups of trials (e.g., novel versus familiar; unpredicted reward versus unpredicted no-reward cue). For each index, we specified the precise temporal window and groups of trials being compared in the Main text or Figure legends. Significance was of these indices was measured with Wilcoxon rank sum tests as previously (68).

Novelty signal latencies were performed similarly as previous studies (68, 118, 119). Briefly for two groups of novelty trials (e.g. NP versus FP) we computed AUC in time, comparing spike density functions (SDFs) across the two conditions. Latency of discrimination was when the AUC was *significantly* greater than chance (0.5 AUC being chance; threshold: >=0.6 for novelty-excited, or =<0.4 for novelty inhibited; p-value threshold: 0.05) for at least 30 milliseconds. This approach was repeated on SDFs triggered on distinct events (object onset, target acquisition time) to verify latency differences were not due to differences in neural dynamics. The key advantage of this approach is it accurately reflects the *relative* differences in latency across different brain regions (118), but like all analyses of latency, it does not index the precise true “time” at which the brain explicitly has access to particular information to guide behavior.

### Electrical stimulation and pharmacological inactivation

Low intensity electrical stimulation (50uA, 400 Hz, 300 ms) was delivered from fractal onset on 50% of trials. The stimulation strength was chosen based on previous studies in monkey (74, 77, 78). The stimulation sites in ZI were determined where significant novelty-excited neurons were found in single-unit recording. The stimulation sites in the subthalamic nucleus and the thalamus were determined to be ~1 or ~2 mm apart from stimulation sites in ZI through the electrode path. We also unilaterally injected GABAa agonist, muscimol (8 μg/μl for monkey R, 4 μg/μl for monkey S) into ZI of monkey R and S. The injection sites were determined where neurons showing a significant novelty-related activity were recorded in single-unit recording. The drug solution was pressure-injected, 0.1 μl per minute at 1-minute interval (0.4 to 0.8 μl in total), using a 10-μl microsyringe (Hamilton) with handmade injectrode. The dose was determined based on previous studies in monkeys (122, 123). During each session, the monkeys first performed 250 trials as a preinjection control. The drug was then injected. 15 minutes after the injection, the monkeys started performing post-injection trials for as long as they could perform. We also conducted sham sessions as control in which exactly the same procedures as the muscimol-injections were performed except for putting injectrode into brain.

## REFERENCES

1. Barto A, Mirolli M, Baldassarre G. Novelty or Surprise? Frontiers in psychology. 2013;4(907). doi: 10.3389/fpsyg.2013.00907.

2. Jaegle A, Mehrpour V, Rust N. Visual novelty, curiosity, and intrinsic reward in machine learning and the brain. Current opinion in neurobiology. 2019;58:167–74.

3. Wang T, Mitchell CJ. Attention and relative novelty in human perceptual learning. Journal of experimental psychology Animal behavior processes. 2011;37(4):436–45. Epub 2011/04/20. doi: 10.1037/a0023104. PubMed PMID: 21500932.

4. Ghazizadeh A, Griggs W, Hikosaka O. Ecological origins of object salience: Reward, uncertainty, aversiveness, and novelty. Frontiers in neuroscience. 2016;10:378.

5. Loewenstein G. The psychology of curiosity: A review and reinterpretation. Psychological bulletin. 1994;116(1):75.

6. Butler RA. Discrimination learning by rhesus monkeys to visual-exploration motivation. Journal of Comparative and Physiological Psychology 1953;46(2):95.

7. Berlyne DE. Novelty and curiosity as determinants of exploratory behaviour. British Journal of Psychology. 1950;41(1):68.

8. Cohen JY, Haesler S, Vong L, Lowell BB, Uchida N. Neuron-type-specific signals for reward and punishment in the ventral tegmental area. Nature. 2012;482(7383):85–8. doi: 10.1038/nature10754. PubMed PMID: 22258508; PubMed Central PMCID: PMC3271183.

9. Schultz W, Dayan P, Montague PR. A neural substrate of prediction and reward. Science. 1997;275(5306):1593–9. Epub 1997/03/14. PubMed PMID: 9054347.

10. Bromberg-Martin ES, Matsumoto M, Hikosaka O. Dopamine in motivational control: rewarding, aversive, and alerting. Neuron. 2010;68(5):815–34. Epub 2010/12/15. doi: 10.1016/j.neuron.2010.11.022. PubMed PMID: 21144997; PubMed Central PMCID: PMC3032992.

11. Lisman JE, Grace AA. The hippocampal-VTA loop: controlling the entry of information into long-term memory. Neuron. 2005;46(5):703–13. Epub 2005/06/01. doi: 10.1016/j.neuron.2005.05.002. PubMed PMID: 15924857.

12. Tapper AR, Molas S. Midbrain circuits of novelty processing. Neurobiology of Learning and Memory. 2020:107323.

13. Duszkiewicz AJ, McNamara CG, Takeuchi T, Genzel L. Novelty and dopaminergic modulation of memory persistence: a tale of two systems. Trends in neurosciences. 2019;42(2):102–14.

14. Bunzeck N, Düzel E. Absolute coding of stimulus novelty in the human substantia nigra/VTA. Neuron. 2006;51(3):369–79.

15. Wittmann BC, Bunzeck N, Dolan RJ, Düzel E. Anticipation of novelty recruits reward system and hippocampus while promoting recollection. Neuroimage. 2007;38(1):194–202.

16. Lak A, Stauffer WR, Schultz W. Dopamine prediction error responses integrate subjective value from different reward dimensions. Proceedings of the National Academy of Sciences of the United States of America. 2014;111(6):2343–8. doi: 10.1073/pnas.1321596111. PubMed PMID: 24453218; PubMed Central PMCID: PMC3926061.

17. Schultz W. Updating dopamine reward signals. Current opinion in neurobiology. 2013;23(2):229–38.

18. Lak A, Stauffer WR, Schultz W. Dopamine neurons learn relative chosen value from probabilistic rewards. eLife. 2016;5:e18044. doi: 10.7554/eLife.18044. PubMed PMID: 27787196.

19. Dabney W, Kurth-Nelson Z, Uchida N, Starkweather CK, Hassabis D, Munos R, Botvinick M. A distributional code for value in dopamine-based reinforcement learning. Nature. 2020:1–5.

20. Bromberg-Martin ES, Hikosaka O. Midbrain dopamine neurons signal preference for advance information about upcoming rewards. Neuron. 2009;63(1):119–26. PubMed PMID: 19607797.

21. !!! INVALID CITATION !!!

22. McHenry JA, Otis JM, Rossi MA, Robinson JE, Kosyk O, Miller NW, McElligott ZA, Budygin EA, Rubinow DR, Stuber GD. Hormonal gain control of a medial preoptic area social reward circuit. Nature neuroscience. 2017;20(3):449–58.

23. Gunaydin LA, Grosenick L, Finkelstein JC, Kauvar IV, Fenno LE, Adhikari A, Lammel S, Mirzabekov JJ, Airan RD, Zalocusky KA. Natural neural projection dynamics underlying social behavior. Cell. 2014;157(7):1535–51.

24. Ranganath C, Rainer G. Neural mechanisms for detecting and remembering novel events. Nature reviews Neuroscience. 2003;4(3):193–202. Epub 2003/03/04. doi: 10.1038/nrn1052. PubMed PMID: 12612632.

25. Menegas W, Babayan BM, Uchida N, Watabe-Uchida M. Opposite initialization to novel cues in dopamine signaling in ventral and posterior striatum in mice. eLife. 2017;6:e21886. doi: 10.7554/eLife.21886.

26. Woloszyn L, Sheinberg DL. Effects of Long-term Visual Experience on Responses of Distinct Classes of Single Units in Inferior Temporal Cortex. Neuron. 2012;74(1):193–205. doi: 10.1016/j.neuron.2012.01.032. PubMed PMID: PMC3329224.

27. Kakade S, Dayan P. Dopamine: generalization and bonuses. Neural Networks. 2002;15(4-6):549–59.

28. Huang X, Weng J. Novelty and reinforcement learning in the value system of developmental robots 2002.

29. Oudeyer P-Y, Kaplan F, Hafner VV. Intrinsic motivation systems for autonomous mental development. IEEE transactions on evolutionary computation. 2007;11(2):265–86.

30. Gottlieb J, Lopes M, Oudeyer P-Y. Motivated cognition: Neural and computational mechanisms of curiosity, attention, and intrinsic motivation. Recent developments in neuroscience research on human motivation: Emerald Group Publishing Limited; 2016. p. 149–72.

31. Monosov IE. How Outcome Uncertainty Mediates Attention, Learning, and Decision-Making. Trends in neurosciences. 2020. doi: https://doi.org/10.1016/j.tins.2020.06.009.

32. Brown MW, Aggleton JP. Recognition memory: what are the roles of the perirhinal cortex and hippocampus? Nature Reviews Neuroscience. 2001;2(1):51–61.

33. Higuchi S-I, Miyashita Y. Formation of mnemonic neuronal responses to visual paired associates in inferotemporal cortex is impaired by perirhinal and entorhinal lesions. Proceedings of the National Academy of Sciences. 1996;93(2):739–43.

34. Haskins AL, Yonelinas AP, Quamme JR, Ranganath C. Perirhinal cortex supports encoding and familiarity-based recognition of novel associations. Neuron. 2008;59(4):554–60.

35. Murray EA, Richmond BJ. Role of perirhinal cortex in object perception, memory, and associations. Current opinion in neurobiology. 2001;11(2):188–93.

36. Lin CS, Nicolelis MA, Schneider JS, Chapin JK, Jr. GABAergic pathway from zona incerta to neocortex: clarification. Science. 1991;251(4998):1162. Epub 1991/03/18. PubMed PMID: 1706534.

37. Lin CS, Nicolelis MA, Schneider JS, Chapin JK. A major direct GABAergic pathway from zona incerta to neocortex. Science. 1990;248(4962):1553–6. Epub 1990/06/22. PubMed PMID: 2360049.

38. Mitrofanis J. Some certainty for the “zone of uncertainty”? Exploring the function of the zona incerta. Neuroscience. 2005;130(1):1–15. doi: https://doi.org/10.1016/j.neuroscience.2004.08.017.

39. Wang X, Chou X-l, Zhang LI, Tao HW. Zona Incerta: An Integrative Node for Global Behavioral Modulation. Trends in neurosciences. 2020;43(2):82–7. doi: https://doi.org/10.1016/j.tins.2019.11.007.

40. Zhao Z-d, Chen Z, Xiang X, Hu M, Xie H, Jia X, Cai F, Cui Y, Chen Z, Qian L. Zona incerta GABAergic neurons integrate prey-related sensory signals and induce an appetitive drive to promote hunting. Nature neuroscience. 2019;22(6):921–32.

41. Tonelli L, Chiaraviglio E. Enhancement of water intake in rats after lidocaine injection in the zona incerta. Brain Res Bull. 1993;31(1-2):1–5. Epub 1993/01/01. PubMed PMID: 8453481.

42. May PJ, Basso MA. Connections between the zona incerta and superior colliculus in the monkey and squirrel. Brain Structure and Function. 2018;223(1):371–90. doi: 10.1007/s00429-017-1503-2.

43. Lovejoy LP, Krauzlis RJ. Inactivation of primate superior colliculus impairs covert selection of signals for perceptual judgments. Nature neuroscience. 2010;13(2):261–6. doi: 10.1038/nn.2470.

44. Krauzlis RJ, Lovejoy LP, Zénon A. Superior Colliculus and Visual Spatial Attention. Annual review of neuroscience. 2013;36:10.1146/annurev-neuro-062012-170249. doi: 10.1146/annurev-neuro-062012-170249. PubMed PMID: PMC3820016.

45. Lauwereyns J, Watanabe K, Coe B, Hikosaka O. A neural correlate of response bias in monkey caudate nucleus. Nature. 2002;418(6896):413–7. doi: 10.1038/nature00892. PubMed PMID: 12140557.

46. Kim HF, Ghazizadeh A, Hikosaka O. Separate groups of dopamine neurons innervate caudate head and tail encoding flexible and stable value memories. Frontiers in neuroanatomy. 2014;8:120. doi: 10.3389/fnana.2014.00120. PubMed PMID: 25400553; PubMed Central PMCID: PMC4214359.

47. Ghazizadeh A, Griggs W, Hikosaka O. Object-finding skill created by repeated reward experience. Journal of vision. 2016;16(10):17. doi: 10.1167/16.10.17. PubMed PMID: PMC5015994.

48. Jutras MJ, Buffalo EA. Recognition memory signals in the macaque hippocampus. Proceedings of the National Academy of Sciences. 2010;107(1):401–6.

49. Zhang K, Chen CD, Monosov IE. Novelty, Salience, and Surprise Timing Are Signaled by Neurons in the Basal Forebrain. Current Biology. 2019;29(1):134–42.e3. doi: https://doi.org/10.1016/j.cub.2018.11.012.

50. Tachibana Y, Hikosaka O. The primate ventral pallidum encodes expected reward value and regulates motor action. Neuron. 2012;76(4):826–37. doi: 10.1016/j.neuron.2012.09.030. PubMed PMID: 23177966; PubMed Central PMCID: PMC3519929.

51. Hikosaka O. Basal ganglia mechanisms of reward-oriented eye movement. Annals of the New York Academy of Sciences. 2007;1104:229–49. doi: 10.1196/annals.1390.012. PubMed PMID: 17360800.

52. Schultz W. Behavioral theories and the neurophysiology of reward. Annu Rev Psychol. 2006;57:87–115. PubMed PMID: 16318590.

53. Hollerman JR, Tremblay L, Schultz W. Influence of reward expectation on behavior-related neuronal activity in primate striatum. Journal of neurophysiology. 1998;80(2):947–63.

54. Matsumoto M, Hikosaka O. Two types of dopamine neuron distinctly convey positive and negative motivational signals. Nature. 2009;459(7248):837–41. Epub 2009/05/19. doi: 10.1038/nature08028. PubMed PMID: 19448610; PubMed Central PMCID: PMC2739096.

55. Watabe-Uchida M, Zhu L, Ogawa SK, Vamanrao A, Uchida N. Whole-brain mapping of direct inputs to midbrain dopamine neurons. Neuron. 2012;74(5):858–73. doi: 10.1016/j.neuron.2012.03.017. PubMed PMID: 22681690.

56. Hollerman JR, Schultz W. Dopamine neurons report an error in the temporal prediction of reward during learning. Nature neuroscience. 1998;1(4):304–9.

57. Joshua M, Adler A, Mitelman R, Vaadia E, Bergman H. Midbrain dopaminergic neurons and striatal cholinergic interneurons encode the difference between reward and aversive events at different epochs of probabilistic classical conditioning trials. The Journal of neuroscience : the official journal of the Society for Neuroscience. 2008;28(45):11673–84. doi: 10.1523/JNEUROSCI.3839-08.2008. PubMed PMID: 18987203.

58. Eshel N, Bukwich M, Rao V, Hemmelder V, Tian J, Uchida N. Arithmetic and local circuitry underlying dopamine prediction errors. Nature. 2015;525(7568):243–6.

59. Hikosaka O. The habenula: from stress evasion to value-based decision-making. Nature reviews Neuroscience. 2010;11(7):503–13. doi: 10.1038/nrn2866. PubMed PMID: 20559337; PubMed Central PMCID: PMC3447364.

60. Matsumoto M, Hikosaka O. Lateral habenula as a source of negative reward signals in dopamine neurons. Nature. 2007;447(7148):1111–5. Epub 2007/05/25. doi: 10.1038/nature05860. PubMed PMID: 17522629.

61. Matsumoto M, Hikosaka O. Negative motivational control of saccadic eye movement by the lateral habenula. Progress in brain research. 2008;171:399–402. Epub 2008/08/23. doi: 10.1016/S0079-6123(08)00658-4. PubMed PMID: 18718332; PubMed Central PMCID: PMC2735791.

62. Matsumoto M, Hikosaka O. Representation of negative motivational value in the primate lateral habenula. Nature neuroscience. 2009;12(1):77–84. Epub 2008/12/02. doi: 10.1038/nn.2233. PubMed PMID: 19043410; PubMed Central PMCID: PMC2737828.

63. Kawai T, Yamada H, Sato N, Takada M, Matsumoto M. Roles of the Lateral Habenula and Anterior Cingulate Cortex in Negative Outcome Monitoring and Behavioral Adjustment in Nonhuman Primates. Neuron. 2015;88(4):792–804. doi: 10.1016/j.neuron.2015.09.030. PubMed PMID: 26481035.

64. Salas R, Baldwin P, De Biasi M, Montague R. BOLD responses to negative reward prediction errors in human habenula. Frontiers in human neuroscience. 2010;4:36.

65. Stephenson-Jones M, Yu K, Ahrens S, Tucciarone JM, van Huijstee AN, Mejia LA, Penzo MA, Tai L-H, Wilbrecht L, Li B. A basal ganglia circuit for evaluating action outcomes. Nature. 2016;539(7628):289–93.

66. Bromberg-Martin ES, Hikosaka O. Lateral habenula neurons signal errors in the prediction of reward information. Nature neuroscience. 2011;14(9):1209–16. doi: 10.1038/nn.2902. PubMed PMID: 21857659; PubMed Central PMCID: PMC3164948.

67. Bromberg-Martin ES, Matsumoto M, Hikosaka O. Distinct tonic and phasic anticipatory activity in lateral habenula and dopamine neurons. Neuron. 2010;67(1):144–55. Epub 2010/07/14. doi: 10.1016/j.neuron.2010.06.016. PubMed PMID: 20624598; PubMed Central PMCID: PMC2905384.

68. White JK, Bromberg-Martin ES, Heilbronner SR, Zhang K, Pai J, Haber SN, Monosov IE. A neural network for information seeking. Nature communications. 2019;10(1):1–19.

69. Ljungberg T, Apicella P, Schultz W. Responses of monkey dopamine neurons during learning of behavioral reactions. Journal of neurophysiology. 1992;67(1):145–63. doi: 10.1152/jn.1992.67.1.145.

70. Morrens J, Aydin Ç, van Rensburg AJ, Rabell JE, Haesler S. Cue-evoked dopamine promotes conditioned responding during learning. Neuron. 2020;106(1):142–53. e7.

71. Costa VD, Tran VL, Turchi J, Averbeck BB. Dopamine modulates novelty seeking behavior during decision making. Behavioral Neuroscience. 2014;128(5):556–66. doi: 10.1037/a0037128.

72. Schultz W. Recent advances in understanding the role of phasic dopamine activity. F1000Research. 2019;8.

73. Monosov IE, Thompson KG. Frontal eye field activity enhances object identification during covert visual search. Journal of neurophysiology. 2009;102(6):3656–72. Epub 2009/10/16. doi: 10.1152/jn.00750.2009. PubMed PMID: 19828723; PubMed Central PMCID: PMC2804410.

74. Yamamoto S, Monosov IE, Yasuda M, Hikosaka O. What and where information in the caudate tail guides saccades to visual objects. The Journal of neuroscience : the official journal of the Society for Neuroscience. 2012;32(32):11005–16. Epub 2012/08/10. doi: 10.1523/JNEUROSCI.0828-12.2012. PubMed PMID: 22875934; PubMed Central PMCID: PMC3465728.

75. McIntyre CC, Grill WM. Selective microstimulation of central nervous system neurons. Annals of biomedical engineering. 2000;28(3):219–33.

76. McIntyre CC, Grill WM. Extracellular stimulation of central neurons: influence of stimulus waveform and frequency on neuronal output. Journal of neurophysiology. 2002;88(4):1592–604.

77. Salzman CD, Newsome WT. Neural mechanisms for forming a perceptual decision. Science. 1994;264(5156):231–7.

78. Ballesta S, Shi W, Conen KE, Padoa-Schioppa C. Values encoded in orbitofrontal cortex are causally related to economic choices. Nature. 2020;588(7838):450–3.

79. Odegaard B, Grimaldi P, Cho SH, Peters MAK, Lau H, Basso MA. Superior colliculus neuronal ensemble activity signals optimal rather than subjective confidence. Proceedings of the National Academy of Sciences. 2018. doi: 10.1073/pnas.1711628115.

80. Trageser JC, Keller A. Reducing the uncertainty: gating of peripheral inputs by zona incerta. Journal of Neuroscience. 2004;24(40):8911–5.

81. Trageser JC, Burke KA, Masri R, Li Y, Sellers L, Keller A. State-dependent gating of sensory inputs by zona incerta. Journal of neurophysiology. 2006;96(3):1456–63.

82. Barthó P, Slézia A, Varga V, Bokor H, Pinault D, Buzsáki G, Acsády L. Cortical control of zona incerta. Journal of Neuroscience. 2007;27(7):1670–81.

83. Xiang J-Z, Brown M. Differential neuronal encoding of novelty, familiarity and recency in regions of the anterior temporal lobe. Neuropharmacology. 1998;37(4-5):657–76.

84. Wan H, Aggleton JP, Brown MW. Different contributions of the hippocampus and perirhinal cortex to recognition memory. Journal of Neuroscience. 1999;19(3):1142–8.

85. Tamura K, Takeda M, Setsuie R, Tsubota T, Hirabayashi T, Miyamoto K, Miyashita Y. Conversion of object identity to object-general semantic value in the primate temporal cortex. Science. 2017;357(6352):687–92.

86. Furtak SC, Wei SM, Agster KL, Burwell RD. Functional neuroanatomy of the parahippocampal region in the rat: the perirhinal and postrhinal cortices. Hippocampus. 2007;17(9):709–22.

87. Tomás Pereira I, Agster KL, Burwell RD. Subcortical connections of the perirhinal, postrhinal, and entorhinal cortices of the rat. I. afferents. Hippocampus. 2016;26(9):1189–212.

88. Agster KL, Tomás Pereira I, Saddoris MP, Burwell RD. Subcortical connections of the perirhinal, postrhinal, and entorhinal cortices of the rat. II. efferents. Hippocampus. 2016;26(9):1213–30.

89. Lawrence BM, White III RL, Snyder LH. Delay-period activity in visual, visuomovement, and movement neurons in the frontal eye field. Journal of neurophysiology. 2005;94(2):1498–508.

90. Sommer MA, Wurtz RH. Composition and topographic organization of signals sent from the frontal eye field to the superior colliculus. Journal of neurophysiology. 2000;83(4):1979–2001.

91. Schall JD. The neural selection and control of saccades by the frontal eye field. Philosophical Transactions of the Royal Society of London Series B: Biological Sciences. 2002;357(1424):1073–82.

92. de Git KCG, Hazelhoff EM, Nota MHC, Schele E, Luijendijk MCM, Dickson SL, van der Plasse G, Adan RAH. Zona incerta neurons projecting to the ventral tegmental area promote action initiation towards feeding. The Journal of Physiology. 2021;599(2):709–24. doi: https://doi.org/10.1113/JP276513.

93. Ossowska K. Zona incerta as a therapeutic target in Parkinson’s disease. Journal of Neurology. 2020;267(3):591–606. doi: 10.1007/s00415-019-09486-8.

94. Langdon AJ, Sharpe MJ, Schoenbaum G, Niv Y. Model-based predictions for dopamine. Current opinion in neurobiology. 2018;49:1–7. doi: https://doi.org/10.1016/j.conb.2017.10.006.

95. Liu Z, Murray EA, Richmond BJ. Learning motivational significance of visual cues for reward schedules requires rhinal cortex. Nature neuroscience. 2000;3(12):1307–15. Epub 2000/12/02. doi: 10.1038/81841. PubMed PMID: 11100152.

96. Ma TP. Saccade-related omnivectoral pause neurons in the primate zona incerta. Neuroreport. 1996;7(15).

97. Miyashita Y. Perirhinal circuits for memory processing. Nature Reviews Neuroscience. 2019;20(10):577–92.

98. Bromberg-Martin ES, Matsumoto M, Nakahara H, Hikosaka O. Multiple timescales of memory in lateral habenula and dopamine neurons. Neuron. 2010;67(3):499–510. Epub 2010/08/11. doi: 10.1016/j.neuron.2010.06.031. PubMed PMID: 20696385; PubMed Central PMCID: PMC2920878.

99. Bromberg-Martin ES, Matsumoto M, Hong S, Hikosaka O. A pallidus-habenula-dopamine pathway signals inferred stimulus values. Journal of neurophysiology. 2010;104(2):1068–76. Epub 2010/06/12. doi: 10.1152/jn.00158.2010. PubMed PMID: 20538770; PubMed Central PMCID: PMC2934919.

100. Nakahara H, Itoh H, Kawagoe R, Takikawa Y, Hikosaka O. Dopamine neurons can represent context-dependent prediction error. Neuron. 2004;41(2):269–80.

101. Morris G, Nevet A, Arkadir D, Vaadia E, Bergman H. Midbrain dopamine neurons encode decisions for future action. Nature neuroscience. 2006;9(8):1057–63.

102. Bromberg-Martin ES, Monosov IE. Neural circuitry of information seeking. Current Opinion in Behavioral Sciences. 2020;35:62–70. doi: https://doi.org/10.1016/j.cobeha.2020.07.006.

103. Ledbetter MN, Chen DC, Monosov IE. Multiple mechanisms for processing reward uncertainty in the primate basal forebrain. The Journal of neuroscience : the official journal of the Society for Neuroscience. 2016;36(30). doi: 10.1523/JNEUROSCI.1123-16.2016.

104. Shin S, Sommer MA. Activity of neurons in monkey globus pallidus during oculomotor behavior compared with that in substantia nigra pars reticulata. Journal of neurophysiology. 2010;103(4):1874–87.

105. Bogadhi AR, Bollimunta A, Leopold DA, Krauzlis RJ. Brain regions modulated during covert visual attention in the macaque. Scientific reports. 2018;8(1):1–15.

106. Paneri S, Gregoriou GG. Top-down control of visual attention by the prefrontal cortex. Functional specialization and long-range interactions. Frontiers in neuroscience. 2017;11:545.

107. Isoda M, Hikosaka O. Role for subthalamic nucleus neurons in switching from automatic to controlled eye movement. The Journal of neuroscience : the official journal of the Society for Neuroscience. 2008;28(28):7209–18. Epub 2008/07/11. doi: 10.1523/JNEUROSCI.0487-08.2008. PubMed PMID: 18614691; PubMed Central PMCID: PMC2667154.

108. Hikosaka O, Wurtz RH. Visual and oculomotor functions of monkey substantia nigra pars reticulata. IV. Relation of substantia nigra to superior colliculus. Journal of neurophysiology. 1983;49(5):1285–301. PubMed PMID: 6306173.

109. Daye PM, Monosov IE, Hikosaka O, Leopold DA, Optican LM. pyElectrode: an open-source tool using structural MRI for electrode positioning and neuron mapping. Journal of neuroscience methods. 2013;213(1):123–31. Epub 2012/12/25. doi: 10.1016/j.jneumeth.2012.12.012. PubMed PMID: 23261658; PubMed Central PMCID: PMC3570753.

110. Dotson NM, Hoffman SJ, Goodell B, Gray CM. A large-scale semi-chronic microdrive recording system for non-human primates. Neuron. 2017;96(4):769–82. e2.

111. Dotson NM, Hoffman SJ, Goodell B, Gray CM. Feature-based visual short-term memory is widely distributed and hierarchically organized. Neuron. 2018;99(1):215–26. e4.

112. Monosov IE, Leopold DA, Hikosaka O. Neurons in the Primate Medial Basal Forebrain Signal Combined Information about Reward Uncertainty, Value, and Punishment Anticipation. The Journal of neuroscience : the official journal of the Society for Neuroscience. 2015;35(19):7443–59. doi: 10.1523/JNEUROSCI.0051-15.2015. PubMed PMID: 25972172; PubMed Central PMCID: PMC4429151.

113. Monosov IE, Hikosaka O. Regionally distinct processing of rewards and punishments by the primate ventromedial prefrontal cortex. J Neurosci. 2012;32(30):10318–30. Epub 2012/07/28. doi: 10.1523/JNEUROSCI.1801-12.2012. PubMed PMID: 22836265; PubMed Central PMCID: PMC3438659.

114. Mao D, Avila E, Caziot B, Laurens J, Dickman JD, Angelaki DE. Spatial representations in macaque hippocampal formation. bioRxiv. 2020:2020.10.03.324848. doi: 10.1101/2020.10.03.324848.

115. Markov NT, Ercsey-Ravasz MM, Ribeiro Gomes A, Lamy C, Magrou L, Vezoli J, Misery P, Falchier A, Quilodran R, Gariel M-A. A weighted and directed interareal connectivity matrix for macaque cerebral cortex. Cerebral cortex. 2014;24(1):17–36.

116. Ogasawara T, Nejime M, Takada M, Matsumoto M. Primate nigrostriatal dopamine system regulates saccadic response inhibition. Neuron. 2018;100(6):1513–26. e4.

117. Ungless MA, Grace AA. Are you or aren’t you? Challenges associated with physiologically identifying dopamine neurons. Trends in neurosciences. 2012;35(7):422–30.

118. Monosov IE, Trageser JC, Thompson KG. Measurements of simultaneously recorded spiking activity and local field potentials suggest that spatial selection emerges in the frontal eye field. Neuron. 2008;57(4):614–25. Epub 2008/02/29. doi: 10.1016/j.neuron.2007.12.030. PubMed PMID: 18304489; PubMed Central PMCID: PMC2350203.

119. Monosov IE, Sheinberg DL, Thompson KG. Paired neuron recordings in the prefrontal and inferotemporal cortices reveal that spatial selection precedes object identification during visual search. Proceedings of the National Academy of Sciences of the United States of America. 2010;107(29):13105–10. Epub 2010/07/10. doi: 10.1073/pnas.1002870107. PubMed PMID: 20615946; PubMed Central PMCID: PMC2919901.

120. Fisher RA. Statistical methods for research workers. Breakthroughs in statistics: Springer; 1992. p. 66–70.

121. Rolls ET, Xiang J-Z. Reward-spatial view representations and learning in the primate hippocampus. Journal of Neuroscience. 2005;25(26):6167–74.

122. Watanabe K, Kimura M. Dopamine receptor-mediated mechanisms involved in the expression of learned activity of primate striatal neurons. Journal of neurophysiology. 1998;79(5):2568–80. doi: 10.1152/jn.1998.79.5.2568. PubMed PMID: 9582229.

123. Sawaguchi T, Goldman-Rakic PS. The role of D1-dopamine receptor in working memory: local injections of dopamine antagonists into the prefrontal cortex of rhesus monkeys performing an oculomotor delayed-response task. Journal of neurophysiology. 1994;71(2):515–28. doi: 10.1152/jn.1994.71.2.515. PubMed PMID: 7909839.

