## Supplemental Figures for "Neuronal mechanisms of novelty seeking"

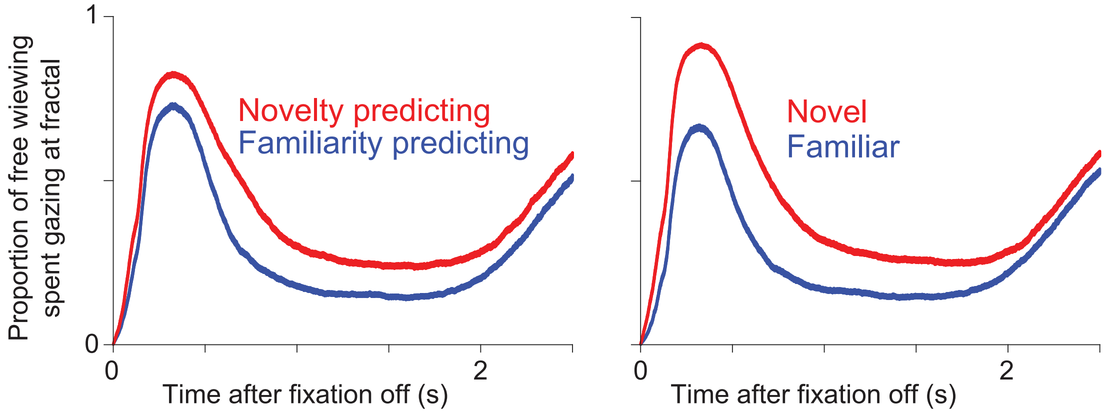


Supplemental Figure 1

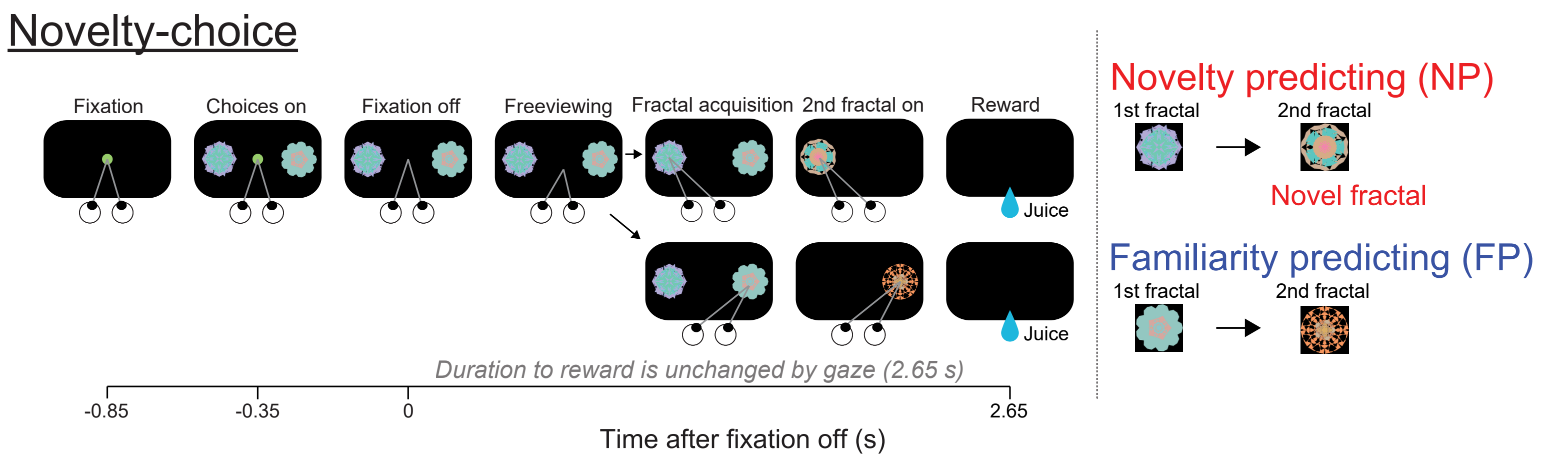


Supplemental Figure 2

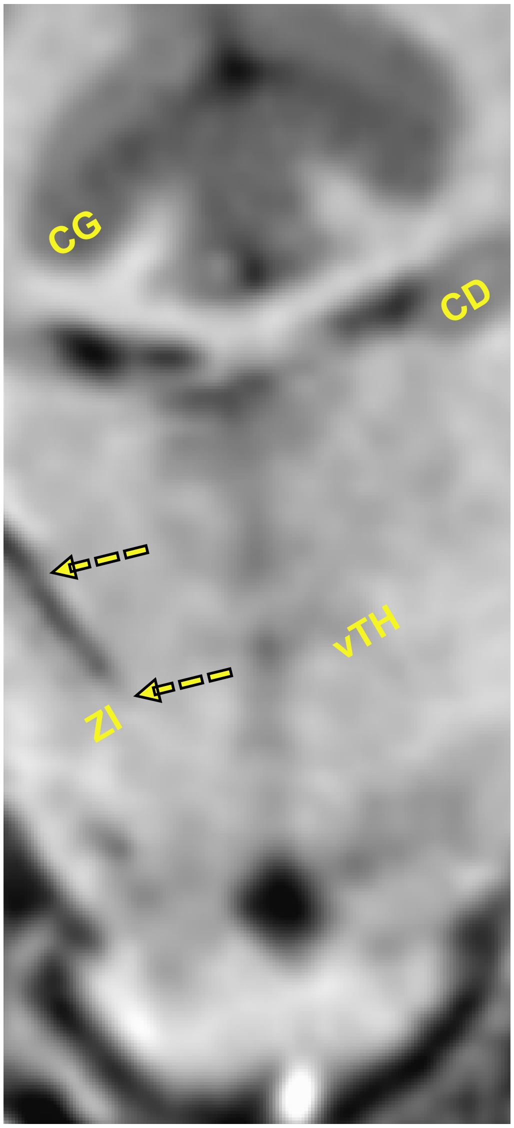


Supplemental Figure 3

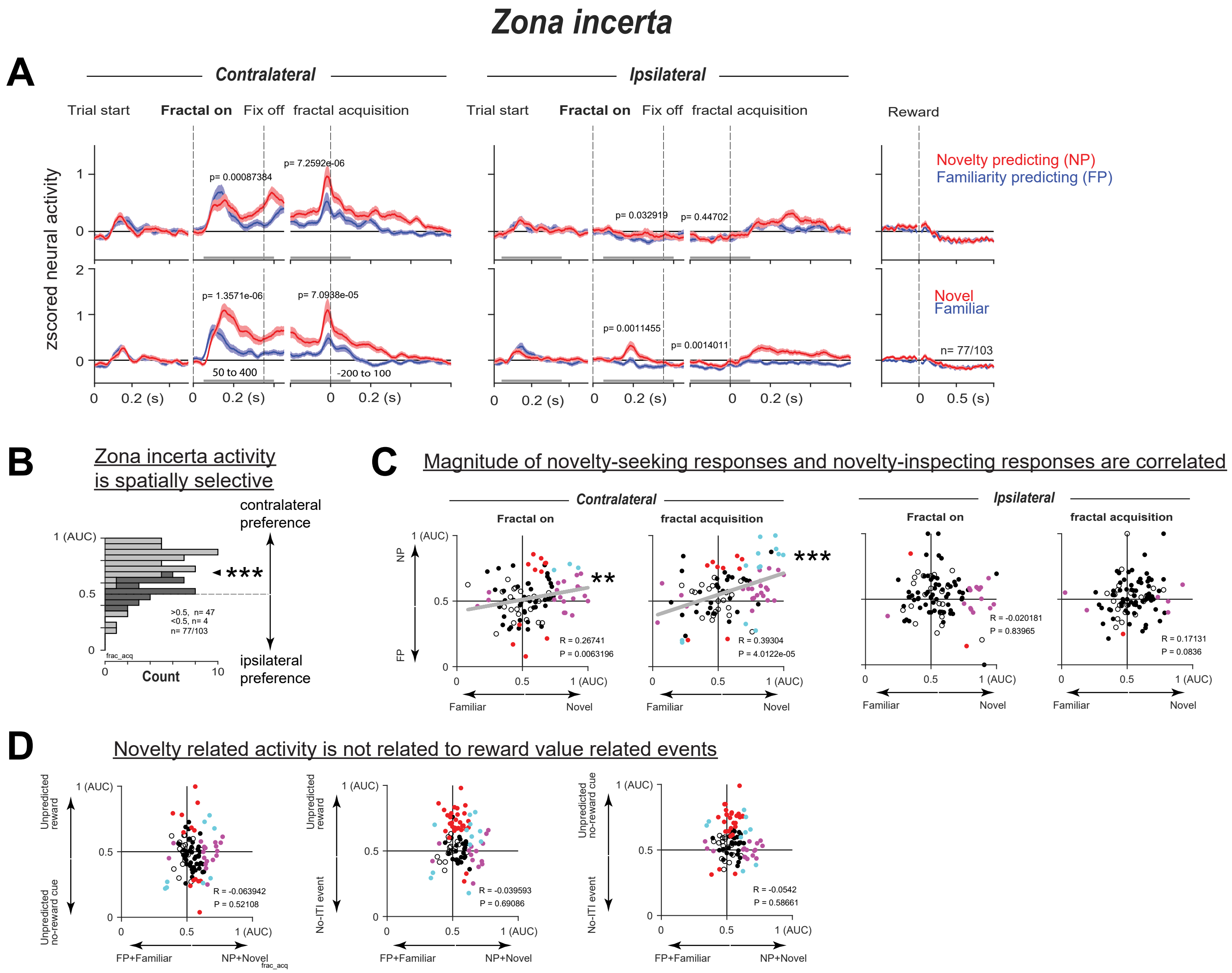


Supplemental Figure 4

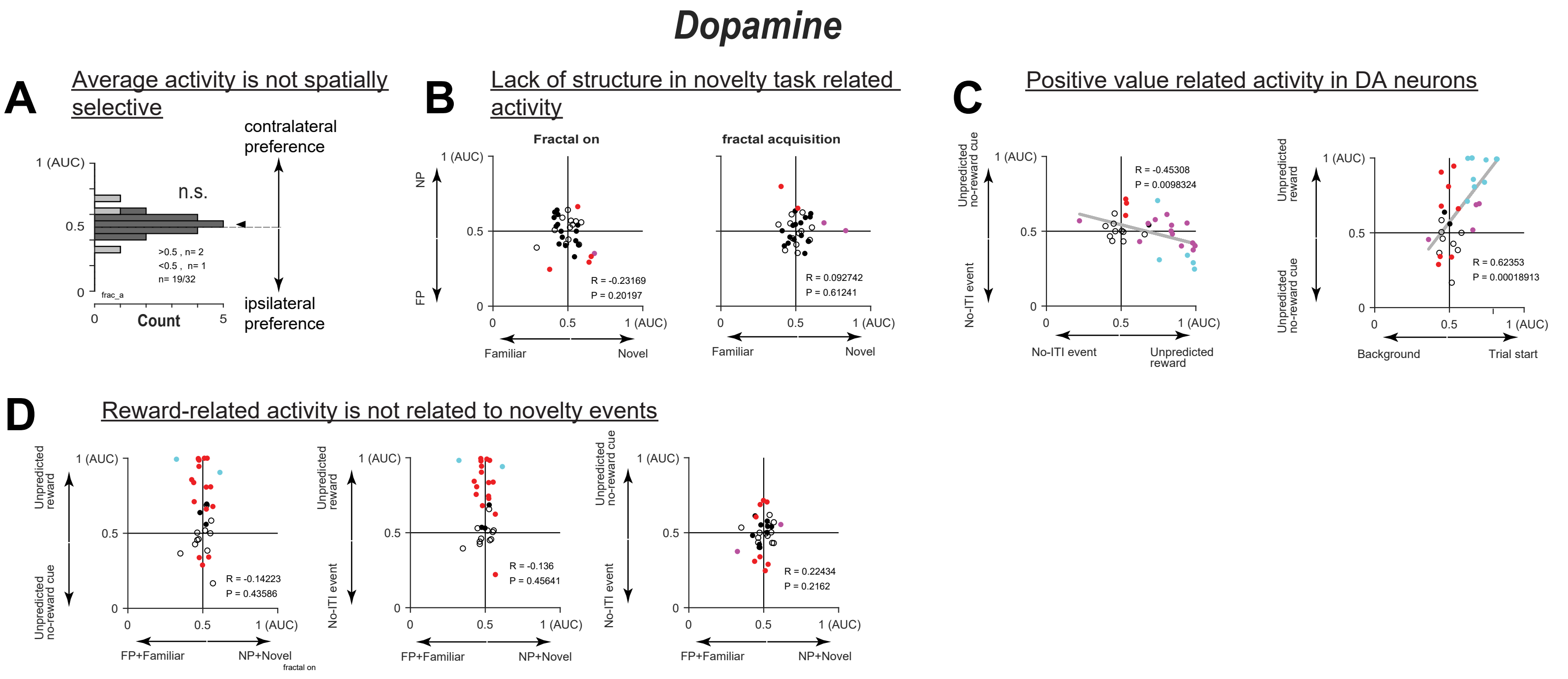


Supplemental Figure 5

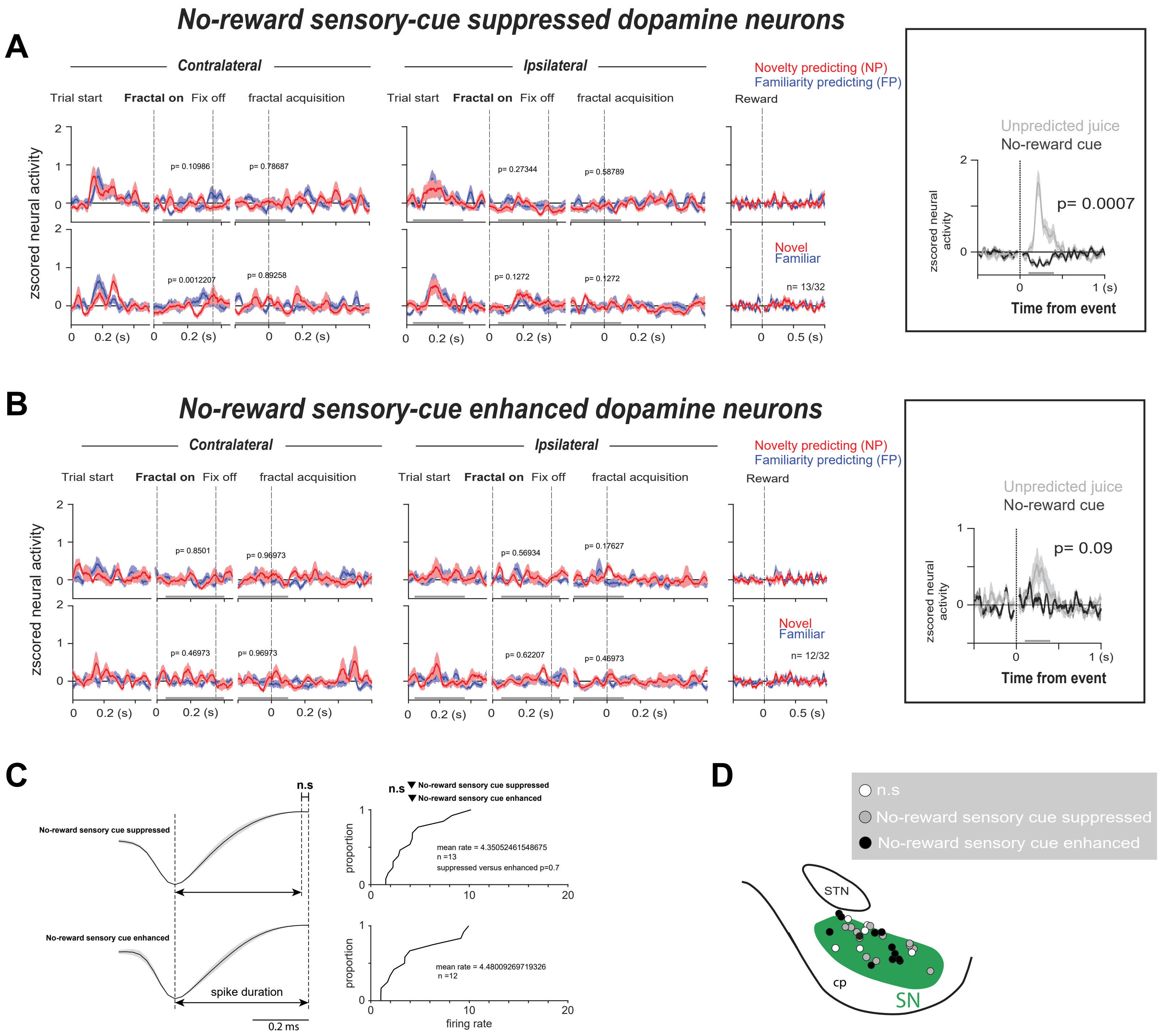


Supplemental Figure 6

### *Habenula*

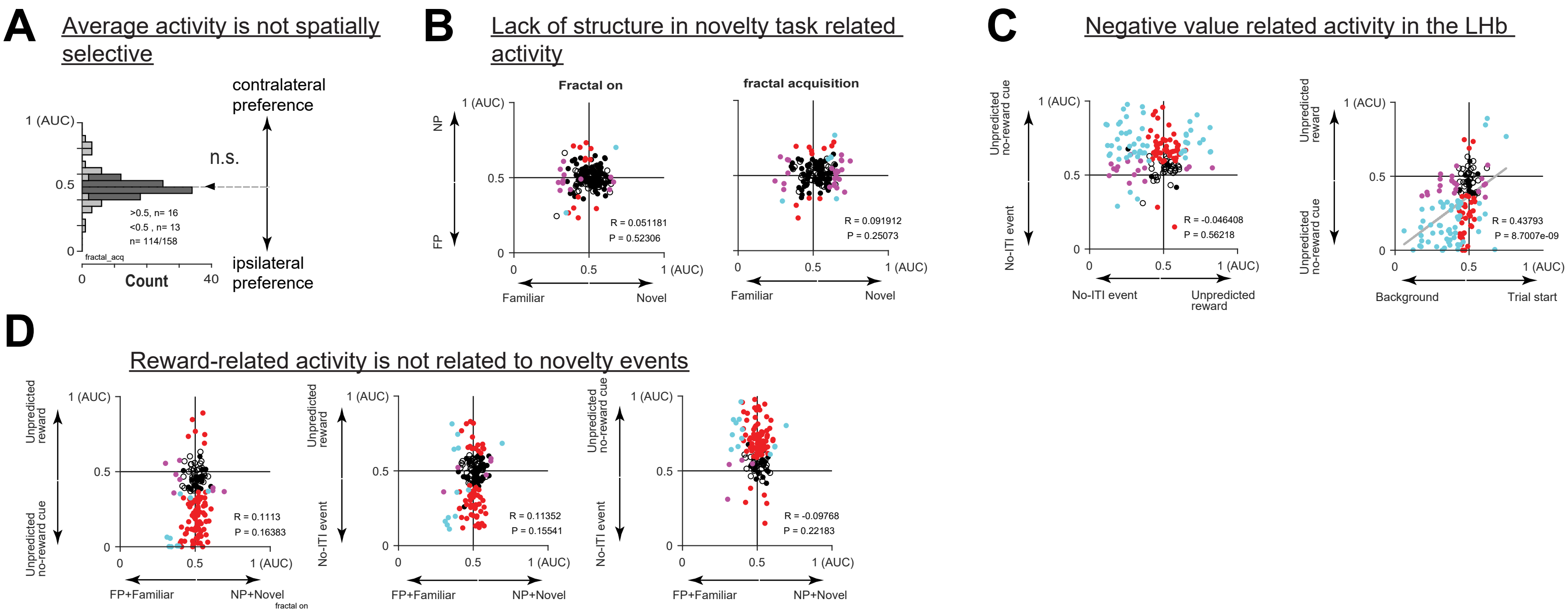


Supplemental Figure 7

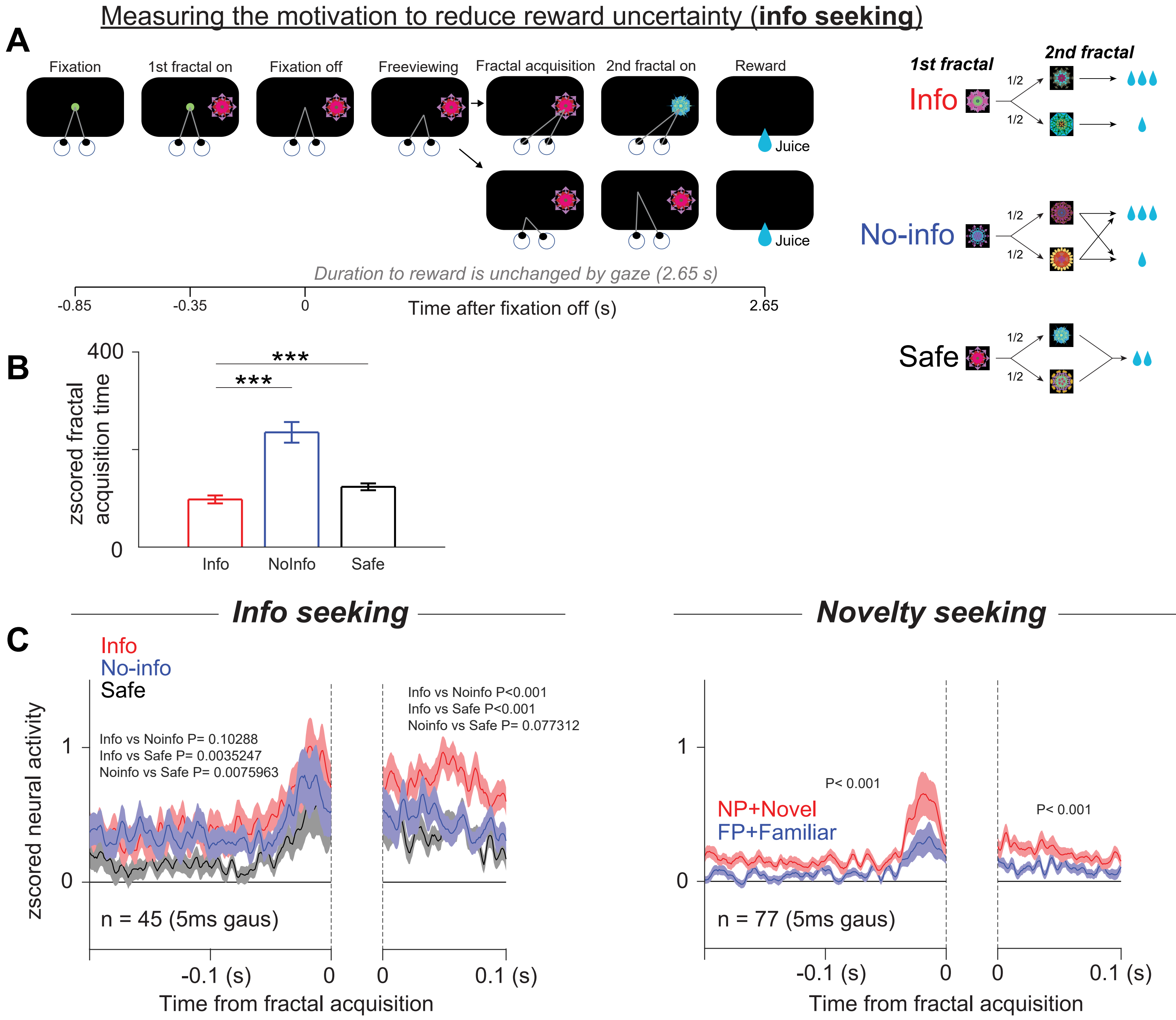


Supplemental Figure 8

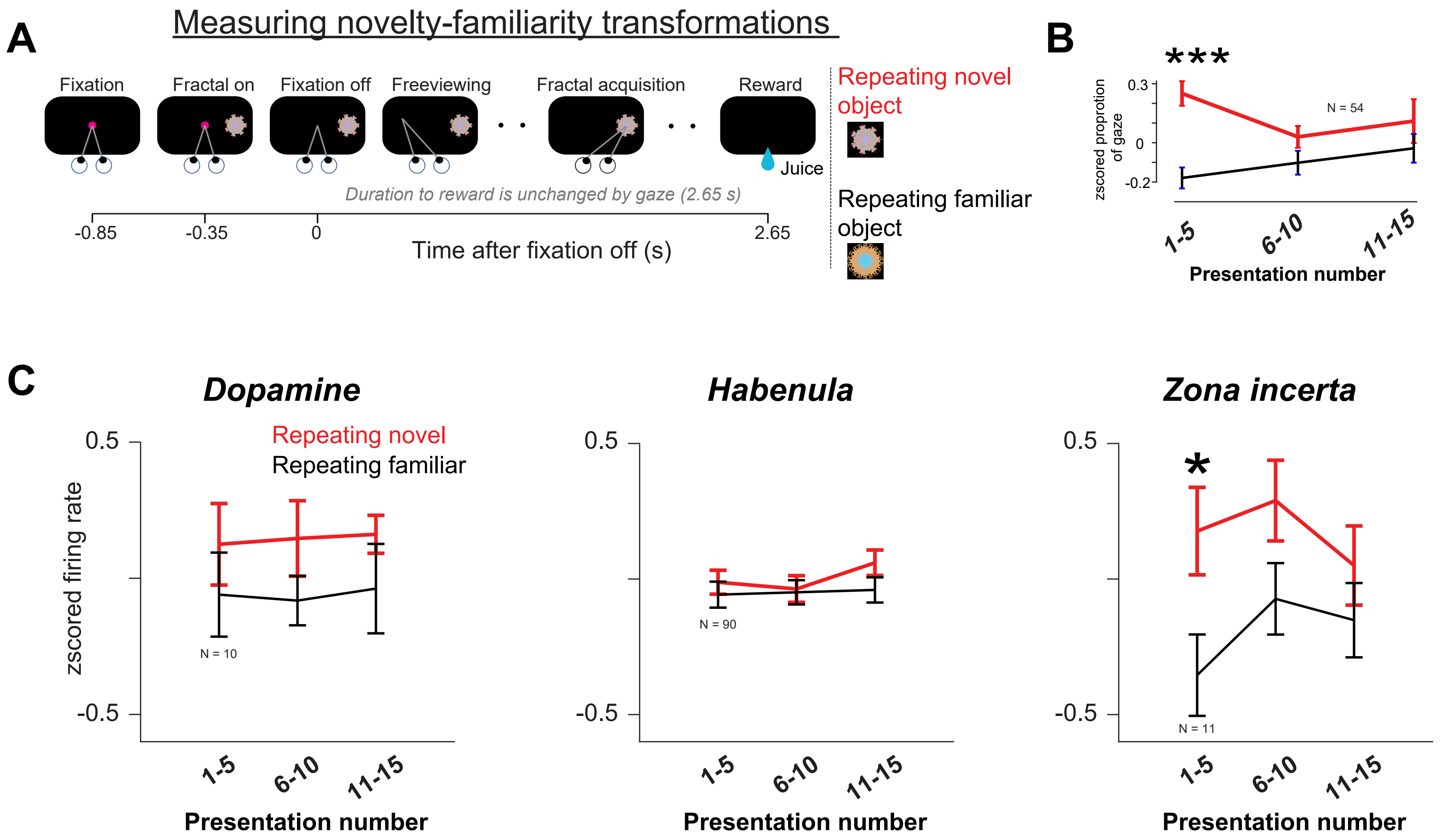


Supplemental Figure 9

### *Basal forebrain*

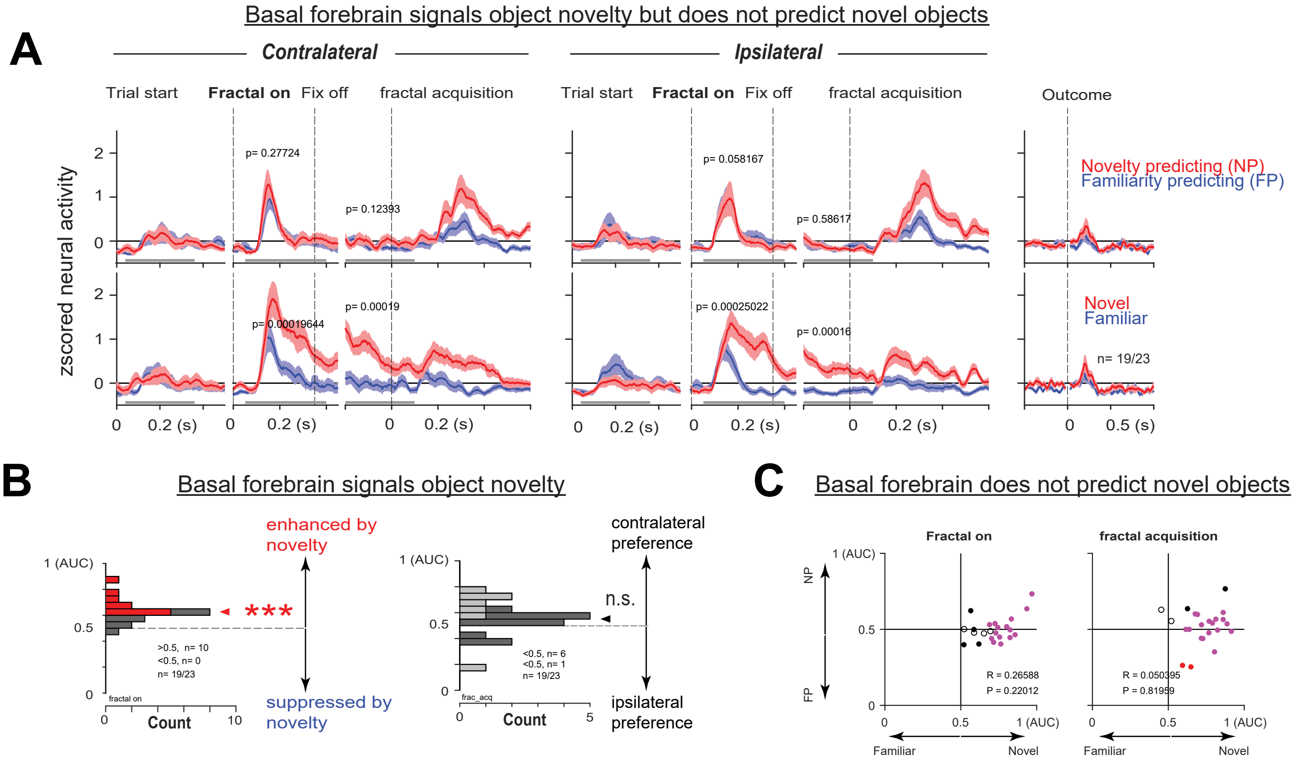


Supplemental Figure 10

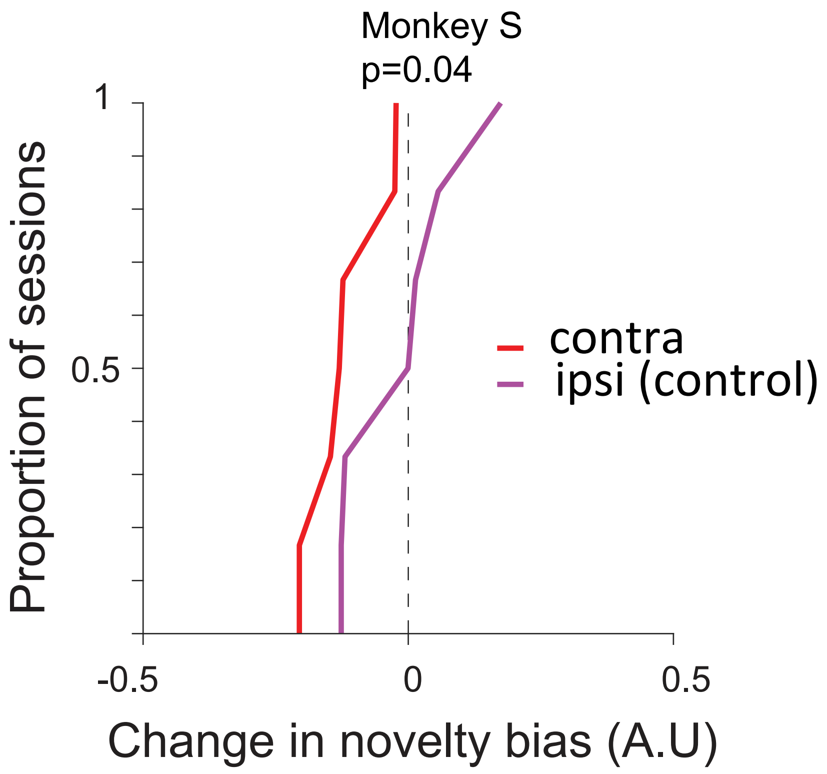


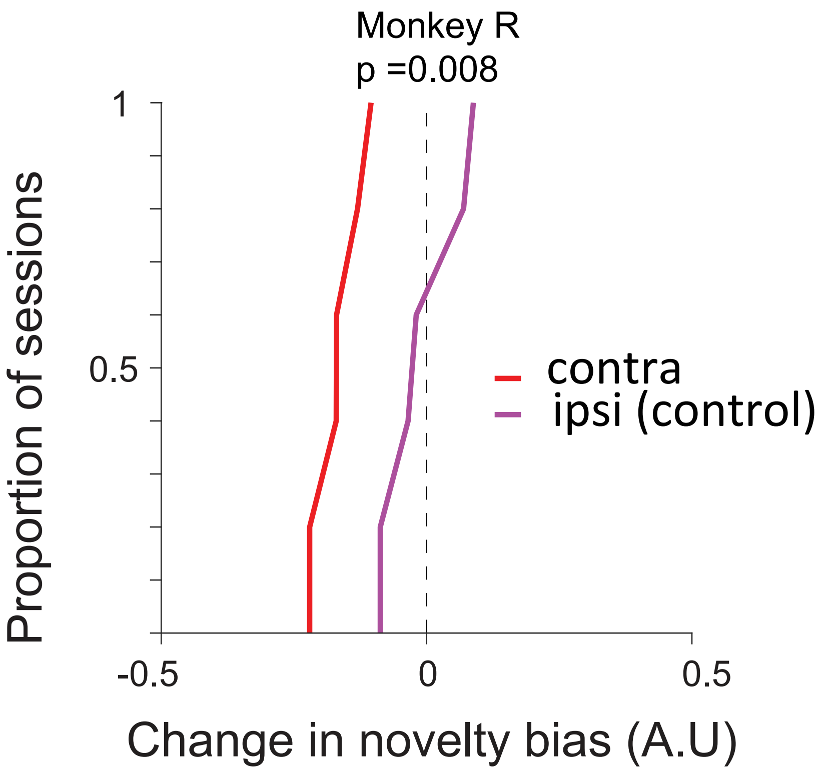


Supplemental Figure 11

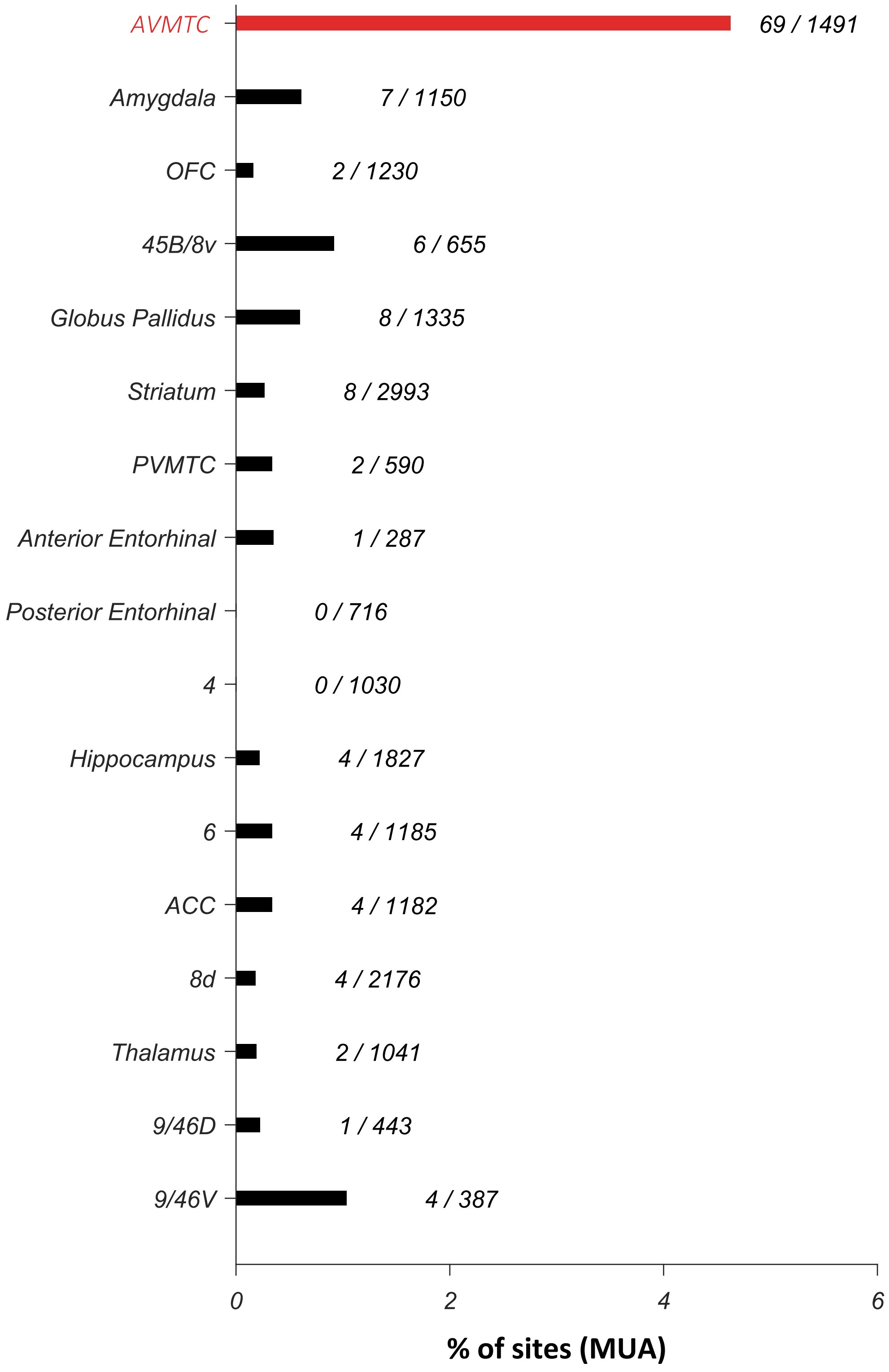


Supplemental Figure 12

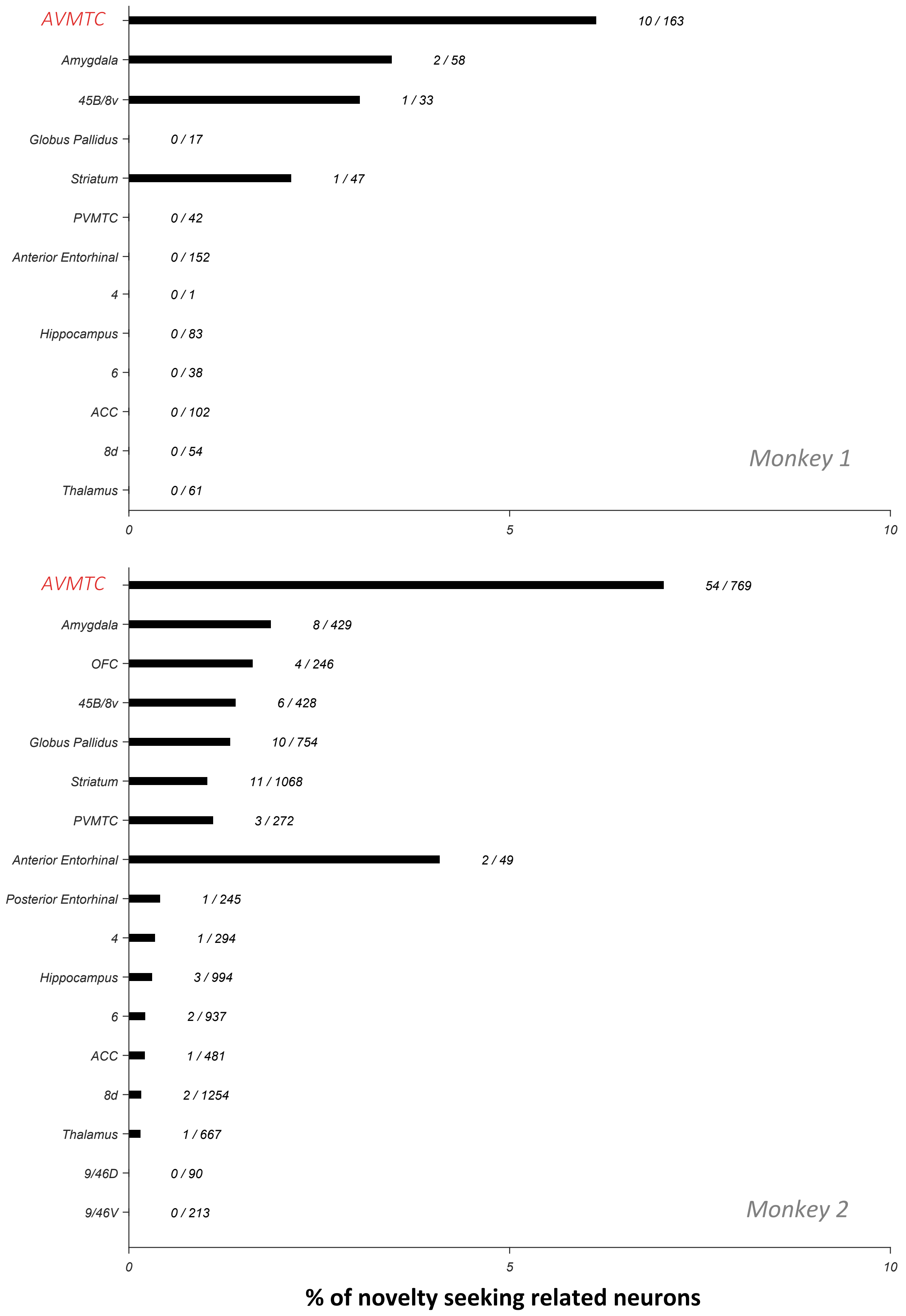


Supplemental Figure 13

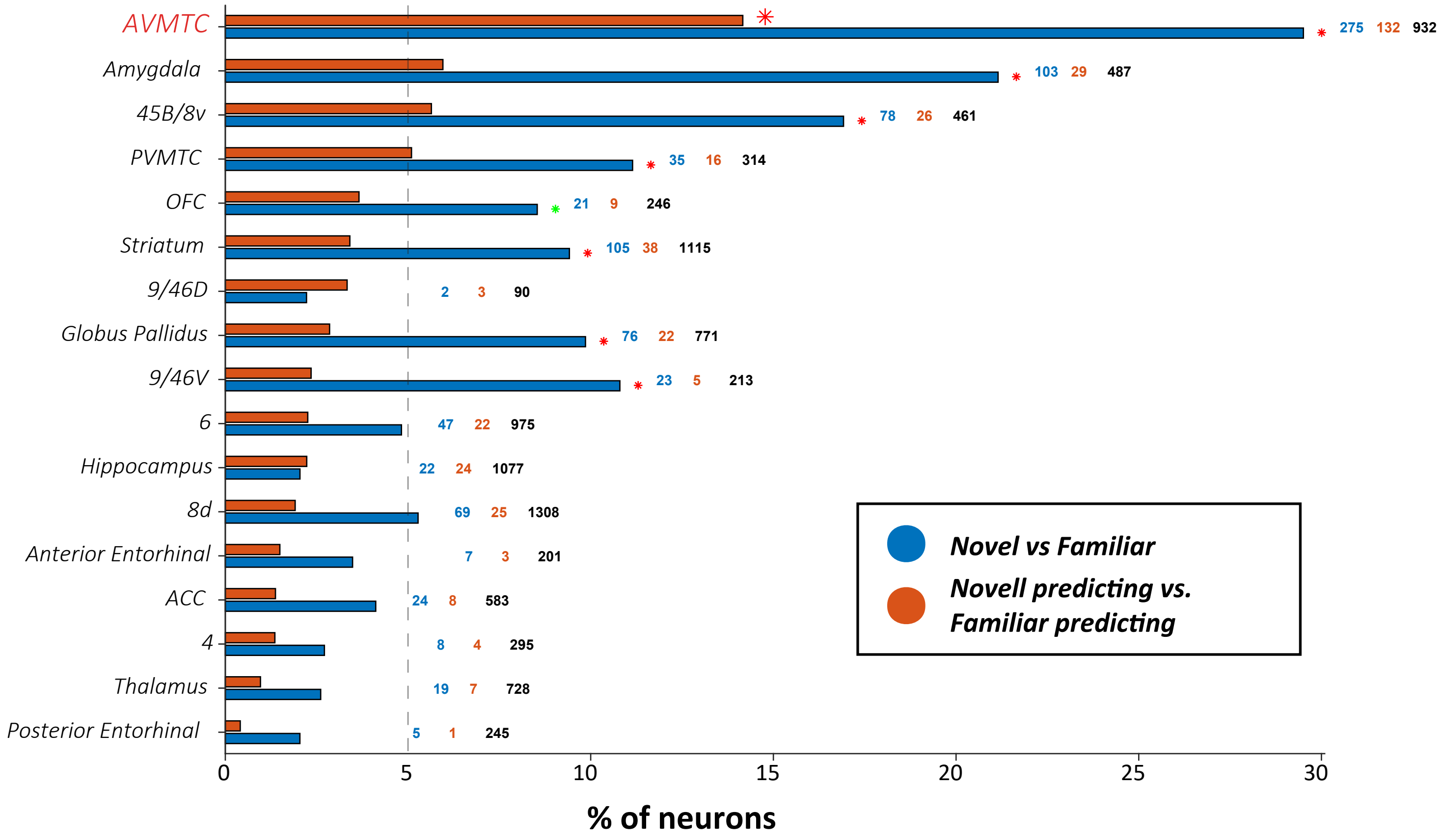


Supplemental Figure 14

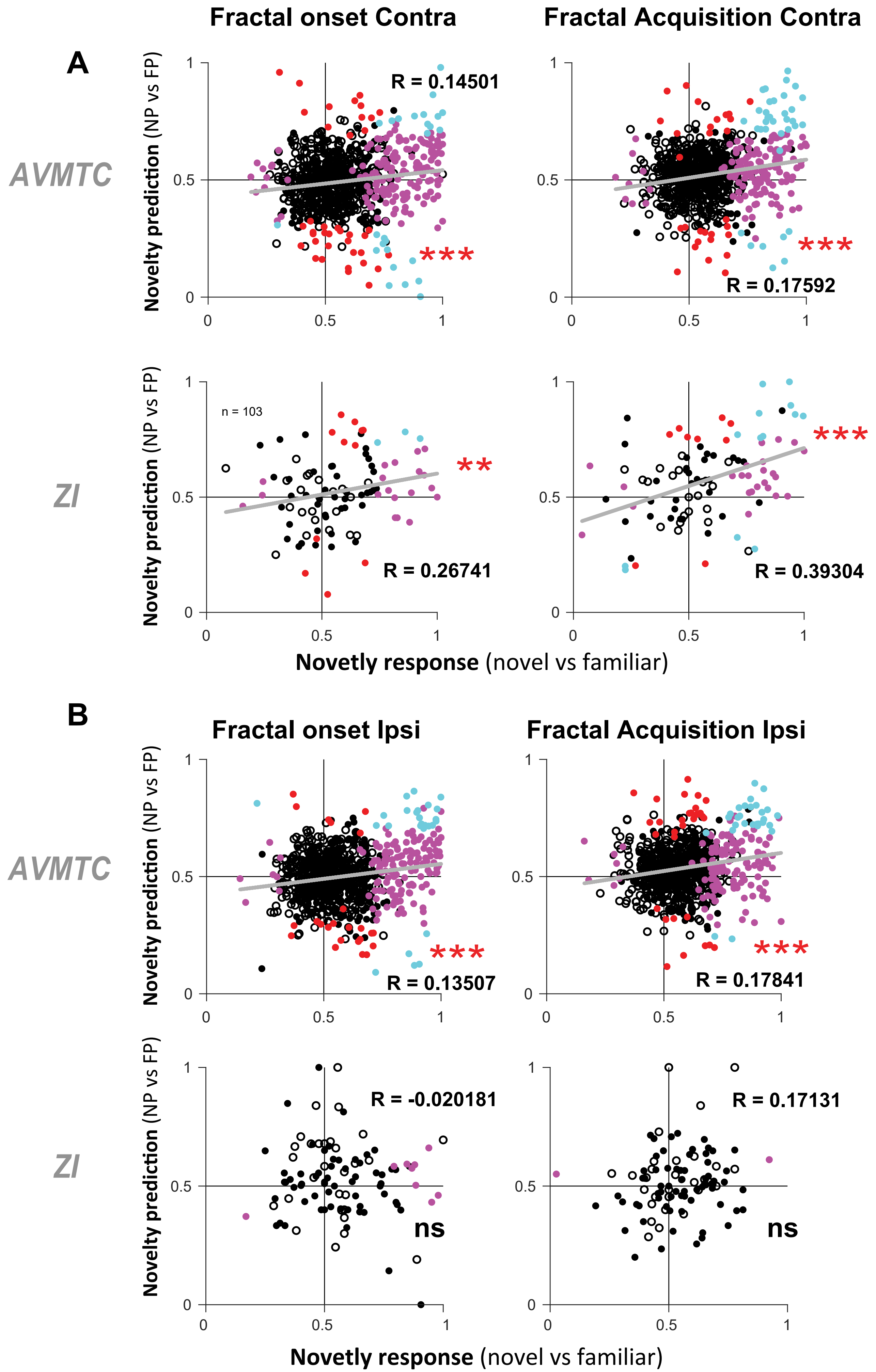


Supplemental Figure 15

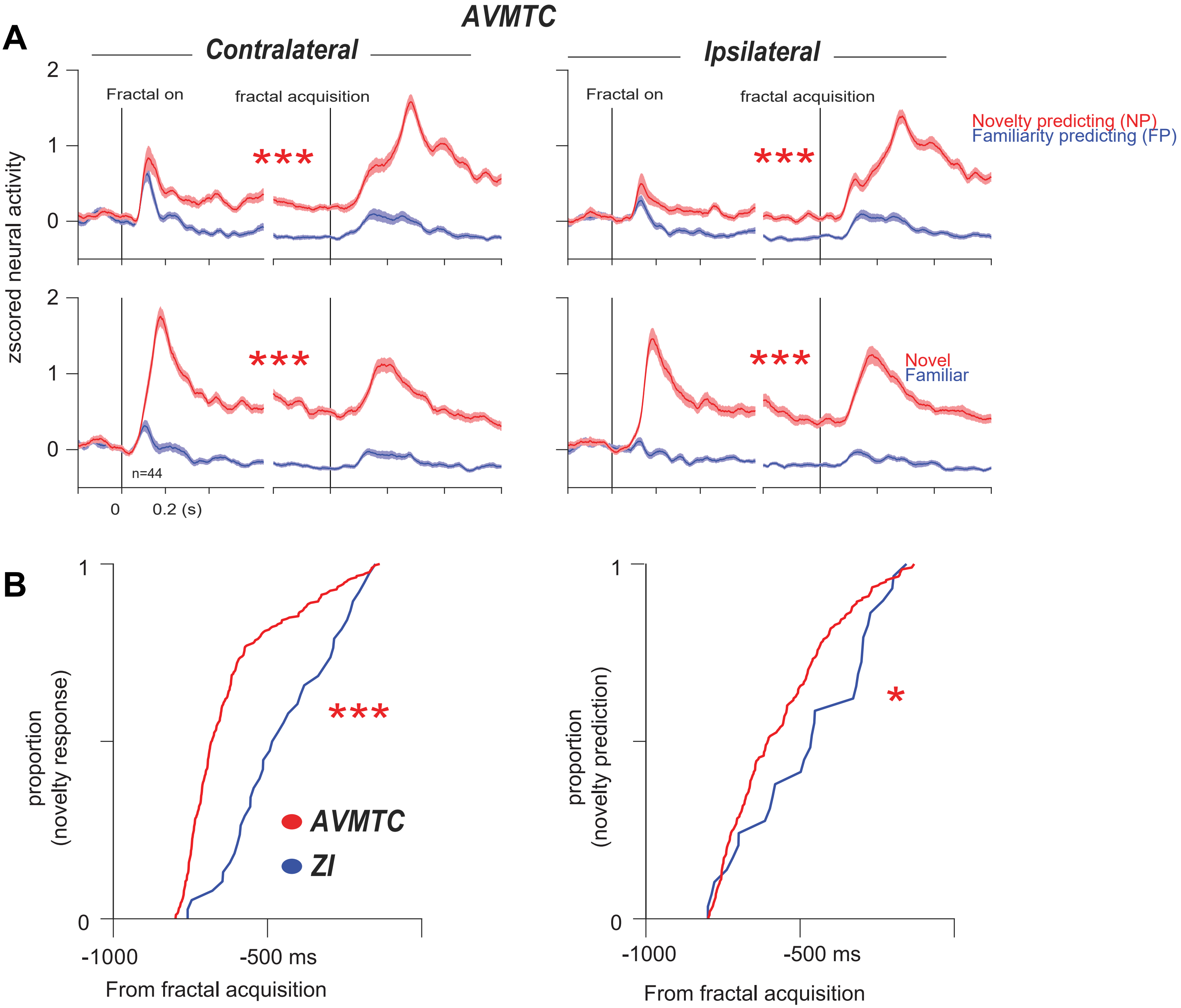


Supplemental Figure 16

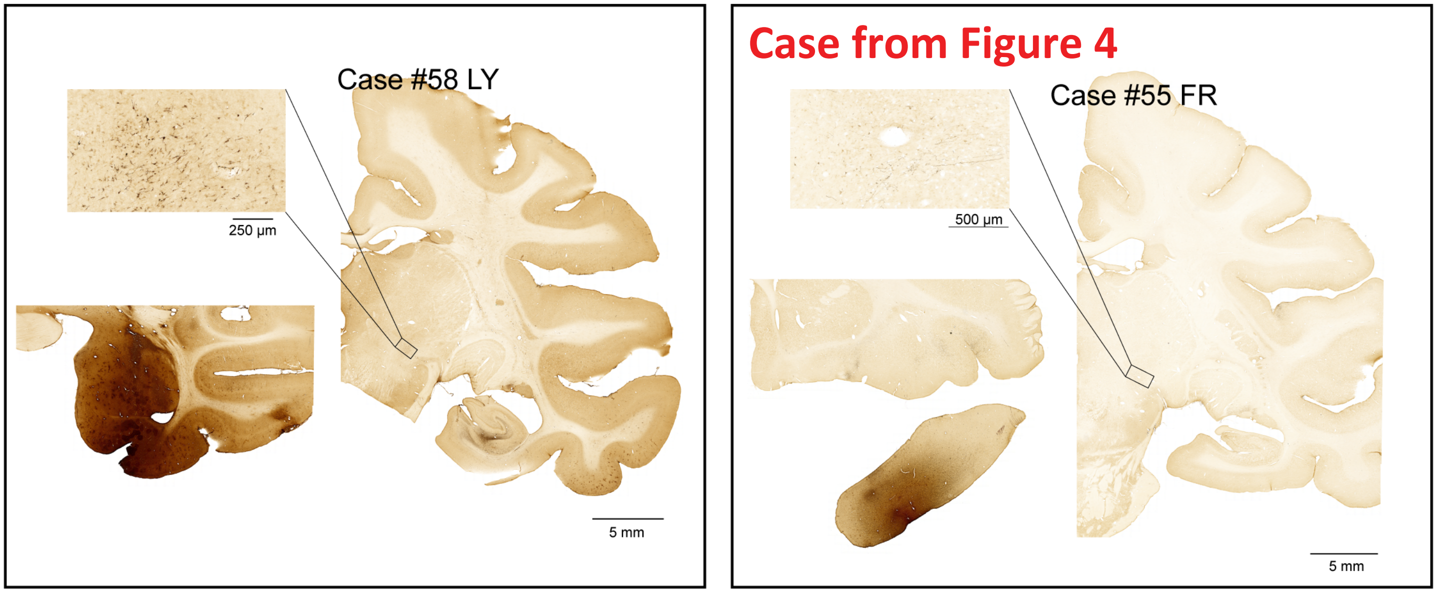


Supplemental Figure 17

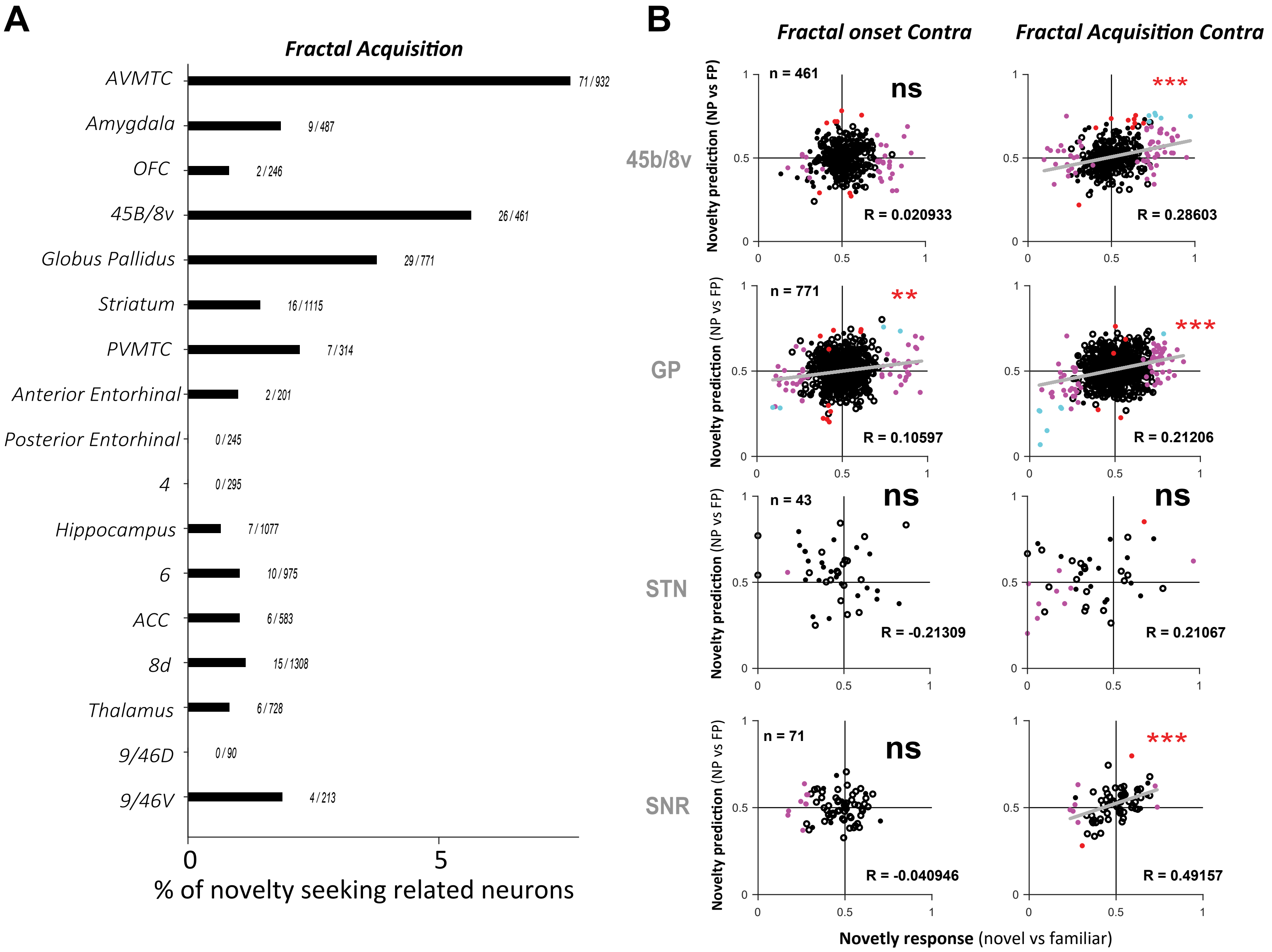


Supplemental Figure 18

# A

| region | cell # | mean(L) - AC(L) | median(L) - AC(L) | std(L) | \|mean(L) - min(L)\| | \|mean(L) - max(L)\| | mean(A) - AC(A) | median(A) - AC(A) | std(A) | \|mean(A) - min(A)\| | \|mean(A) - max(A)\| | mean(D) - AC(D) | median(D) - AC(D) | std(D) | \|mean(D) - min(D)\| | \|mean(D) - max(D)\| |
|---|---|---|---|---|---|---|---|---|---|---|---|---|---|---|---|---|
| AVMTC | 163 | 12.5 | 11.8 | 1.89 | 2.19 | 3.91 | -0.922 | -1.38 | 1.74 | 3.51 | 2.59 | -17.6 | -17.6 | 1.15 | 1.23 | 2.87 |
| Amygdala | 58 | 10.6 | 13.4 | 3 | 3.39 | 2.71 | -0.561 | 1.67 | 2.56 | 3.86 | 2.23 | -9.58 | -10.7 | 1.57 | 2.56 | 3.44 |
| 45B/8v | 33 | 11.8 | 11.8 | 3.61e-15 | 3.55e-15 | 3.55e-15 | 4.73 | 4.73 | 3.61e-15 | 3.55e-15 | 3.55e-15 | 8.11 | 8.11 | 1.8e-15 | 1.78e-15 | 1.78e-15 |
| Globus Pallidus | 17 | 5.99 | 5.72 | 0.599 | 0.269 | 1.25 | -1.1 | -1.37 | 1.11 | 1.79 | 1.26 | 0.0755 | 0.211 | 0.383 | 1.11 | 0.386 |
| Striatum | 47 | 8.41 | 8.75 | 2.77 | 2.7 | 6.46 | -1.95 | -2.89 | 5.01 | 10.1 | 5.15 | 2.79 | -0.413 | 5.46 | 6.56 | 7.97 |
| PVMTC | 42 | 14.9 | 14.9 | 0.0017 | 0.00214 | 0.00145 | -7.91 | -5.95 | 2.29 | 2.61 | 1.96 | -16.4 | -17 | 0.807 | 0.686 | 1.1 |
| Anterior Entorhinal | 152 | 8.82 | 8.78 | 1.35 | 1.56 | 3.02 | -2.63 | -2.9 | 1.44 | 1.8 | 2.78 | -16.8 | -16.5 | 0.773 | 1.63 | 1.99 |
| 4 | 1 | 8.74 | 8.74 | 0 | 0 | 0 | -5.93 | -5.93 | 0 | 0 | 0 | 19.8 | 19.8 | 0 | 0 | 0 |
| Hippocampus | 83 | 12.7 | 13.4 | 1.35 | 6.97 | 0.663 | -3.91 | -2.9 | 2.2 | 11.2 | 1.01 | -13.7 | -13.7 | 1.42 | 1.46 | 5.94 |
| 6 | 38 | 6.82 | 7.23 | 0.687 | 1.14 | 0.405 | 0.323 | 1.68 | 2.42 | 4.73 | 1.36 | 15.2 | 12.1 | 5.4 | 3.14 | 9.95 |
| ACC | 102 | 5.7 | 5.7 | 0.00179 | 0.00391 | 0.00196 | -5.34 | -5.94 | 7.4 | 9.74 | 8.55 | 13.4 | 13.7 | 2.15 | 3.32 | 4.48 |
| 8d | 54 | 9.74 | 8.75 | 1.15 | 0.99 | 2.06 | 4.48 | 4.73 | 0.574 | 1.27 | 0.255 | 12.3 | 13.2 | 1.71 | 2.73 | 1.63 |
| Thalamus | 61 | 6.69 | 7.24 | 0.924 | 0.975 | 2.07 | -10.2 | -8.99 | 1.98 | 3.32 | 1.25 | -1.6 | -1.78 | 0.86 | 1.09 | 1.38 |

| region | cell # | mean(L) - AC(L) | median(L) - AC(L) | std(L) | \|mean(L) - min(L)\| | \|mean(L) - max(L)\| | mean(A) - AC(A) | median(A) - AC(A) | std(A) | \|mean(A) - min(A)\| | \|mean(A) - max(A)\| | mean(D) - AC(D) | median(D) - AC(D) | std(D) | \|mean(D) - min(D)\| | \|mean(D) - max(D)\| |
|---|---|---|---|---|---|---|---|---|---|---|---|---|---|---|---|---|
| AVMTC | 769 | 14.6 | 14.7 | 1.56 | 2.93 | 1.96 | -1 | -1.23 | 1.75 | 2.02 | 6.07 | -17.3 | -17.3 | 1.46 | 2.34 | 7.16 |
| Amygdala | 429 | 12.4 | 13 | 1.75 | 5.66 | 2.48 | -0.534 | -1.17 | 1.49 | 2.59 | 3.64 | -11.6 | -11.7 | 1.69 | 4.66 | 5.35 |
| OFC | 246 | 11.8 | 12 | 2.23 | 4.65 | 3.09 | 9.25 | 9.59 | 1.03 | 3.22 | 1.55 | 1.23 | 1.61 | 1.68 | 5.6 | 2.88 |
| 45B/8v | 428 | 14 | 14 | 1.39 | 4.42 | 1.54 | 3.13 | 3.59 | 1.24 | 3.03 | 2.13 | 8.83 | 8.88 | 1.27 | 3.3 | 3.96 |
| Globus Pallidus | 754 | 9.05 | 8.66 | 1.99 | 3.89 | 6.49 | -3.34 | -3.18 | 2.18 | 6.85 | 4.7 | -1.6 | -1.7 | 1.31 | 3.54 | 3.35 |
| Striatum | 1068 | 11.8 | 13.4 | 3.61 | 7.42 | 5.01 | -3.76 | -5.49 | 5.09 | 6.67 | 11.4 | -1.79 | -3.55 | 5.32 | 7.76 | 10.6 |
| PVMTC | 272 | 15.9 | 15.4 | 0.765 | 0.782 | 1.36 | -5.71 | -5.6 | 2.11 | 6.08 | 1.65 | -17.2 | -17.5 | 1.28 | 1.8 | 4.26 |
| Anterior Entorhinal | 49 | 10 | 10.3 | 0.702 | 1.46 | 0.399 | -2.08 | -1.69 | 1.09 | 1.12 | 1.78 | -17.7 | -17.8 | 0.699 | 0.59 | 1.61 |
| Posterior Entorhinal | 245 | 11.8 | 12.4 | 0.923 | 3.63 | 2.51 | -7.97 | -7.58 | 1.49 | 4.55 | 2.07 | -14.8 | -15.8 | 2.05 | 1.59 | 5.63 |
| 4 | 294 | 9.3 | 9.14 | 3.04 | 3.45 | 6.93 | -8.75 | -9.08 | 1.6 | 3.37 | 4.47 | 17.2 | 16.9 | 1.26 | 5.92 | 4.82 |
| Hippocampus | 994 | 13.7 | 14.2 | 2.15 | 6.07 | 3.47 | -6.98 | -7.23 | 2.85 | 4.93 | 4.46 | -12.1 | -12.2 | 1.96 | 3.69 | 6.92 |
| 6 | 937 | 8.68 | 9.19 | 2.24 | 3.78 | 7.4 | 1.6 | 1.68 | 2.71 | 6.29 | 5.84 | 16.2 | 16.5 | 1.18 | 3.7 | 2.77 |
| ACC | 481 | 5.11 | 5.17 | 1.05 | 1.06 | 1.98 | 7.63 | 8.77 | 3.55 | 18.6 | 2.95 | 13.2 | 13.1 | 1.14 | 2.58 | 2.99 |
| 8d | 1254 | 10.2 | 10.4 | 2.45 | 4.28 | 5.79 | 4.43 | 4.84 | 2.44 | 6.71 | 6.48 | 14.1 | 14 | 1.17 | 3.39 | 2.89 |
| Thalamus | 667 | 7.66 | 7.74 | 1.31 | 1.98 | 3.33 | -9.48 | -9.41 | 1.99 | 2.99 | 4.49 | 2.73 | 2.31 | 2.01 | 6.16 | 4.25 |
| 9/46D | 90 | 12.1 | 11.7 | 1.01 | 2.09 | 1.35 | 9.15 | 9.51 | 1.11 | 2.49 | 1.9 | 12.3 | 13 | 1.58 | 3.37 | 2.5 |
| 9/46V | 213 | 12.8 | 13.1 | 0.616 | 1.19 | 0.351 | 10.9 | 11.2 | 0.59 | 1.41 | 0.271 | 8.73 | 8.55 | 0.554 | 0.614 | 1.35 |

# B

| region | cell # | mean(L) - AC(L) | median(L) - AC(L) | std(L) | \|mean(L) - min(L)\| | \|mean(L) - max(L)\| | mean(A) - AC(A) | median(A) - AC(A) | std(A) | \|mean(A) - min(A)\| | \|mean(A) - max(A)\| | mean(D) - AC(D) | median(D) - AC(D) | std(D) | \|mean(D) - min(D)\| | \|mean(D) - max(D)\| |
|---|---|---|---|---|---|---|---|---|---|---|---|---|---|---|---|---|
| AVMTC | 64 | 14.9 | 15 | 1.86 | 4.62 | 1.59 | -1.42 | -1.23 | 1.46 | 1.48 | 3.4 | -18 | -18.2 | 1.09 | 1.61 | 3.2 |
| Amygdala | 10 | 13 | 13.1 | 1.43 | 1.63 | 1.88 | 0.0541 | 0.153 | 1.56 | 2.72 | 1.62 | -10.4 | -10.4 | 1.14 | 1.87 | 2.69 |
| OFC | 4 | 8.69 | 8.69 | 0.00266 | 0.0023 | 0.0023 | 10.6 | 10.6 | 0.00137 | 0.00118 | 0.00118 | 0.87 | 0.87 | 0.144 | 0.125 | 0.125 |
| 45B/8v | 7 | 13.2 | 13.9 | 1.91 | 3.61 | 2.18 | 2.84 | 3.59 | 1.54 | 2.75 | 1.89 | 8.94 | 8.63 | 1.02 | 1.22 | 1.85 |
| Globus Pallidus | 10 | 10.1 | 8.77 | 3.47 | 3.16 | 5.43 | -5.65 | -4.79 | 3.15 | 4.54 | 3.82 | -1.79 | -1.16 | 2.29 | 3.35 | 2.77 |
| Striatum | 12 | 13 | 13.8 | 3.04 | 6.62 | 3.68 | -5.81 | -7.32 | 4.73 | 4.36 | 10 | -3.98 | -3.93 | 2.23 | 4.67 | 4.32 |
| PVMTC | 3 | 16.1 | 16.6 | 0.866 | 1 | 0.5 | -4.13 | -4.08 | 0.0945 | 0.109 | 0.0546 | -17.1 | -16.8 | 0.484 | 0.559 | 0.279 |
| Anterior Entorhinal | 2 | 8.59 | 8.59 | 0 | 0 | 0 | -0.306 | -0.306 | 0 | 0 | 0 | -16.3 | -16.3 | 0 | 0 | 0 |
| Posterior Entorhinal | 1 | 12.4 | 12.4 | 0 | 0 | 0 | -7.58 | -7.58 | 0 | 0 | 0 | -16.4 | -16.4 | 0 | 0 | 0 |
| 4 | 1 | 11.6 | 11.6 | 0 | 0 | 0 | -5.75 | -5.75 | 0 | 0 | 0 | 16.4 | 16.4 | 0 | 0 | 0 |
| Hippocampus | 3 | 14.9 | 14.3 | 1.27 | 0.802 | 1.47 | -8.28 | -10.4 | 5.04 | 3.63 | 5.76 | -11.4 | -11.4 | 1.04 | 1.01 | 1.06 |
| 6 | 2 | 9.36 | 9.36 | 0.0244 | 0.0173 | 0.0173 | 1.67 | 1.67 | 0.0126 | 0.00888 | 0.00888 | 15.6 | 15.6 | 1.33 | 0.937 | 0.937 |
| ACC | 1 | 5.62 | 5.62 | 0 | 0 | 0 | 8.9 | 8.9 | 0 | 0 | 0 | 11.7 | 11.7 | 0 | 0 | 0 |
| 8d | 2 | 12.4 | 12.4 | 0 | 0 | 0 | 1.96 | 1.96 | 0 | 0 | 0 | 13.6 | 13.6 | 0 | 0 | 0 |
| Thalamus | 1 | 7.82 | 7.82 | 0 | 0 | 0 | -10.9 | -10.9 | 0 | 0 | 0 | 2.11 | 2.11 | 0 | 0 | 0 |

| region | cell # | mean(L) - AC(L) | median(L) - AC(L) | std(L) | \|mean(L) - min(L)\| | \|mean(L) - max(L)\| | mean(A) - AC(A) | median(A) - AC(A) | std(A) | \|mean(A) - min(A)\| | \|mean(A) - max(A)\| | mean(D) - AC(D) | median(D) - AC(D) | std(D) | \|mean(D) - min(D)\| | \|mean(D) - max(D)\| |
|---|---|---|---|---|---|---|---|---|---|---|---|---|---|---|---|---|
| AVMTC | 71 | 14.5 | 15 | 1.9 | 4.22 | 1.99 | -1.1 | -1.07 | 1.82 | 3.32 | 5.98 | -17.6 | -18.1 | 1.16 | 1.98 | 2.61 |
| Amygdala | 9 | 13.4 | 13.4 | 1.01 | 1.91 | 1.45 | 0.206 | 0.214 | 1.77 | 2.87 | 1.47 | -10.6 | -10.7 | 0.253 | 0.152 | 0.666 |
| OFC | 2 | 11.8 | 11.8 | 2.26 | 1.6 | 1.6 | 10.2 | 10.2 | 0.853 | 0.603 | 0.603 | 2.02 | 2.02 | 0.141 | 0.0995 | 0.0995 |
| 45B/8v | 26 | 13.2 | 13.9 | 1.53 | 3.6 | 2.2 | 3.09 | 3.59 | 1.31 | 2.99 | 1.64 | 8.96 | 8.88 | 0.829 | 1.99 | 1.83 |
| Globus Pallidus | 29 | 9.49 | 8.76 | 2.72 | 2.71 | 6.05 | -4.81 | -4.71 | 2.75 | 5.38 | 4.49 | -2.1 | -1.89 | 1.46 | 3.04 | 2 |
| Striatum | 16 | 11.4 | 12.8 | 3.73 | 6.55 | 4.08 | -5.56 | -7.35 | 4.66 | 4.62 | 8.78 | -0.666 | -2.89 | 6.65 | 7.96 | 10 |
| PVMTC | 7 | 16 | 16.6 | 0.839 | 1.11 | 0.671 | -5.27 | -4.09 | 2.4 | 5.26 | 1.19 | -17.1 | -17.4 | 0.86 | 0.655 | 1.83 |
| Anterior Entorhinal | 2 | 8.02 | 8.02 | 1.08 | 0.762 | 0.762 | -2.9 | -2.9 | 2.16 | 1.52 | 1.52 | -16.6 | -16.6 | 0.413 | 0.292 | 0.292 |
| Hippocampus | 7 | 15.1 | 15.8 | 2.19 | 4.58 | 1.9 | -8.99 | -10.3 | 3.4 | 2.92 | 4.98 | -11.5 | -12.1 | 1.21 | 1.58 | 1.3 |
| 6 | 10 | 8.12 | 8.06 | 1.28 | 1.98 | 1.26 | 1.7 | 1.67 | 1.79 | 4.72 | 1.34 | 16.3 | 16.8 | 1.23 | 1.66 | 1.3 |
| ACC | 6 | 4.97 | 4.92 | 0.921 | 0.873 | 1.11 | 1.52 | 7.99 | 10.7 | 15.1 | 7.38 | 12 | 11.7 | 1.42 | 1.92 | 2.18 |
| 8d | 15 | 10.4 | 10.4 | 2.03 | 3.16 | 5.26 | 4.6 | 4.73 | 1.86 | 3.86 | 2.98 | 13.8 | 13.9 | 1.69 | 3.08 | 2.7 |
| Thalamus | 6 | 7.08 | 6.48 | 1.65 | 1.37 | 2.26 | -9.79 | -8.99 | 1.31 | 2.25 | 0.801 | -1.15 | -1.78 | 1.56 | 1.25 | 3.08 |
| 9/46V | 4 | 13.1 | 13.1 | 0.00115 | 0.000576 | 0.00173 | 11.2 | 11.2 | 0.000592 | 0.000888 | 0.000296 | 8.21 | 8.24 | 0.0625 | 0.0937 | 0.0312 |

Supplemental Figure 19

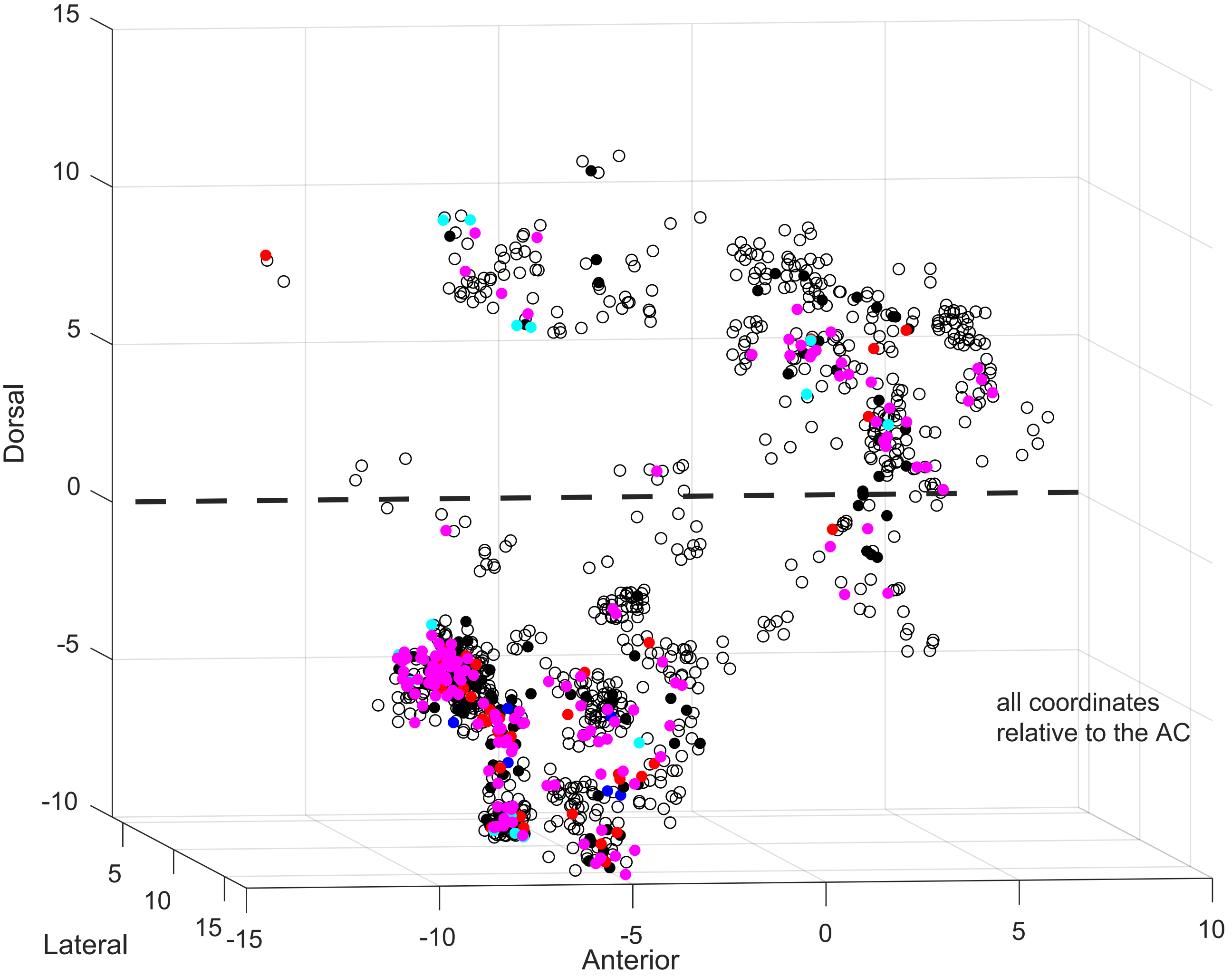


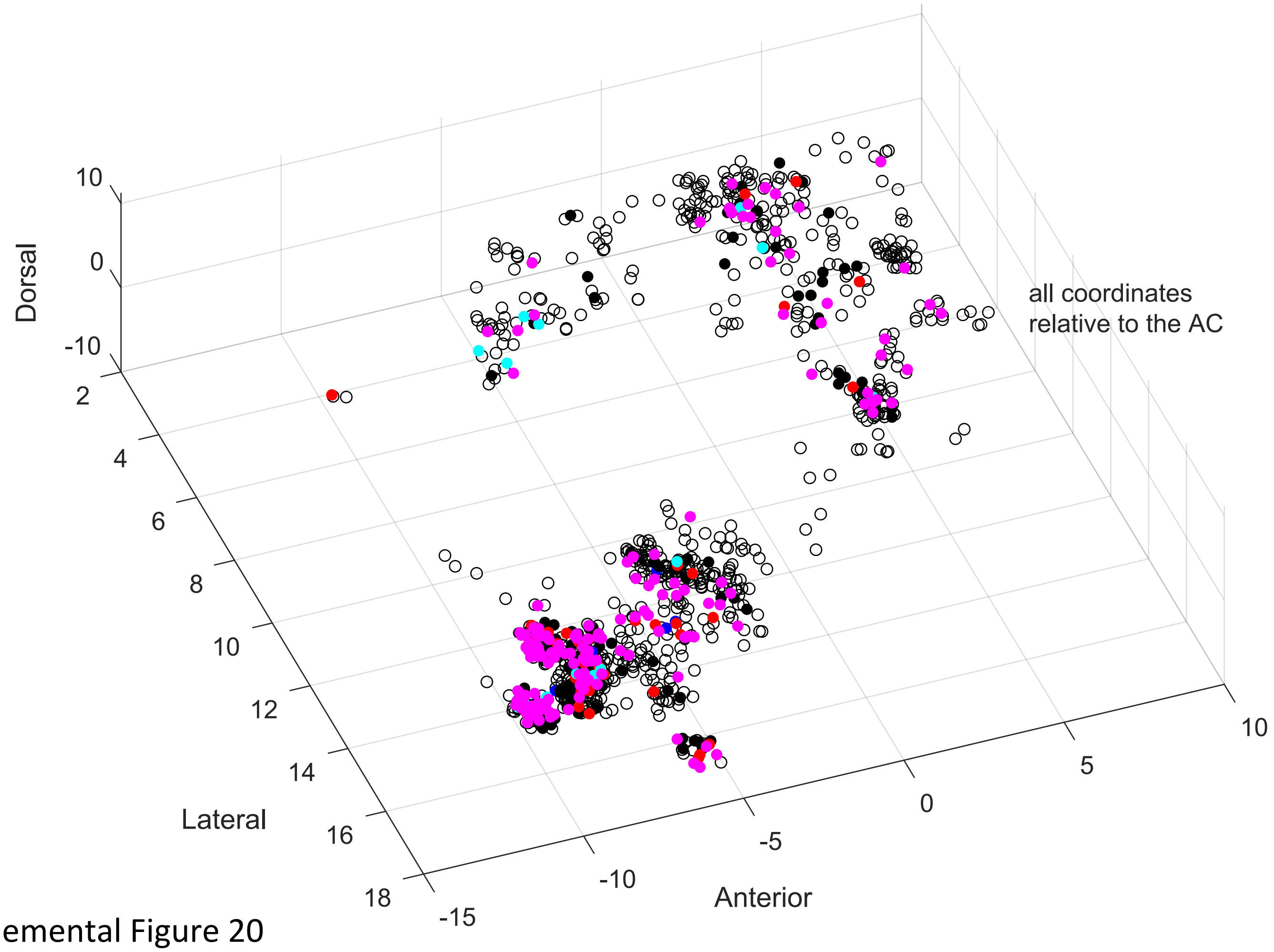


Supplemental Figure 20
